# Metabolite co-variation networks reveal keystone functions and an emergent pathogen state in the human urobiome

**DOI:** 10.64898/2026.08.29.748013

**Authors:** Larissa Della Vedova, Adam J. Bindas, Mariana Teixeira Dias, Jolanda K. Brons, Zizhuang Fang, Aline Marie Fernandes, Pablo Gallardo Molina, David Giron-Villalobos, Thomas Hackl, Jeroen Jansen, Jerry M. Wells, Marjon G.J. de Vos, Celia R. Berkers, Justin J.J van der Hooft

## Abstract

Microbial communities are dynamic, adaptive ecosystems whose collective behavior emerges from metabolic interactions such as cross-feeding, competition, and cooperation, rather than taxonomic diversity or individual metabolic potential alone. This distinction is clinically significant in the postmenopausal urinary tract, where recurrent urinary tract infections (rUTIs) are associated with complex, persistent infection dynamics including multiple contributing bacterial species. The ability of resident microbial communities to prevent pathogen establishment, known as colonization resistance, is increasingly attributed to the metabolic interactions within the urobiome itself rather than any single resident species. However, current approaches, such as taxonomic profiling and classical differential abundance analysis, can only partially describe the presence or maintenance of such interactions. Consequently, the community-level metabolic architecture determining pathogen resistance remains incompletely understood.

To address this gap, we developed PhenoRewire, a network-based framework that quantifies how metabolite co-variation is rewired between biological states using untargeted metabolomics data. We applied this framework to an induced pluripotent stem cell (iPSC) urothelial organoid-derived barrier co-cultured with synthetic urobiome communities as a model of urobiome-pathogen dynamics relevant to rUTIs in two approaches. In an infection model, clinically isolated uropathogens *Escherichia coli* and *Enterococcus faecalis,* were co-cultured with a three-member urobiome community consisting of *Lactobacillus gasseri*, *Lactobacillus crispatus*, and *Gardnerella vaginalis*. Here we show how *E. coli* drove the metabolic reorganization, while *E. faecalis* amplified it disproportionately. PhenoRewire disentangled the 6-fold metabolic network amplification mediated by *E. faecalis* as a metabolic facilitator, revealing an emergent urobiome-pathogen co-variation architecture (1,781 vs 227 edges) not recapitulated by either community alone. Moreover, in a six-member urobiome single-strain dropout experiment, we revealed that removal of the sole Actinomycete *Winkia anitrata* caused significant network collapse (Louvain modularity falls from 0.707 to 0.038), identifying it as the single non-redundant keystone of the community.

More broadly, these results demonstrate how untargeted metabolomics co-variation network analysis can be applied to defined synthetic urobiomes in combination with a urothelial host model to elucidate community dynamics. This framework provides a template that can be extended beyond the urobiome to investigate any complex microbial community where ecological behavior remains an open question.

## Introduction

Microbial communities form coexisting multispecies assemblages shaped by continuous intra-and interspecies interactions that depend on their shared environment. The functional output of such a network can be beneficial, detrimental, or neutral for one or more partners, spanning from mutualism to competition [1]. Diverse mechanisms, including antimicrobial compound secretion, metabolite cross-feeding, and biofilm formation underpin these interactions [2]. Indeed, over the past decades, microbiome research has broadened beyond the description of individual microorganisms inhabiting a given body site or environment to encompass the communities they form and the ecological context in which they exist and interact [3], predominantly through studying their taxonomic composition. However, as metabolically active ecosystems, microbial communities cannot be understood solely from their composition. Both local, pairwise metabolic exchange and indirect, system-level consequences of microbial interactions that propagate across the community can shape ecosystem function, but often remain understudied [4].

The urinary tract microbiome exemplifies this gap. It is now recognized as a metabolically active microbial ecosystem [5–7], yet remains studied predominantly through taxonomic composition. Traditionally, a urinary tract infection (UTI) has been defined by the presence of clinically significant bacteriuria alone, based on the detection of uropathogens like uropathogenic *Escherichia coli* (UPEC) above established quantitative thresholds [9]. While taxonomic profiling provides additional insight into clinical risk and the presence of community members associated with colonization resistance, it does not capture the functional activity or the biological state of the microbial community (i.e., healthy or in dysbiosis) [8].

Capturing these communities therefore requires treating them as dynamic systems whose regulatory state shifts during transitions such as pathogen invasion or keystone loss, rather than as fixed catalogs of taxa. These states typically differ in microbial composition and abundance, shaping community metabolic output. Through competition for nutrients and niches plus the production of inhibitory metabolites, a resident microbial community can suppress the establishment of incoming pathogens, a property termed colonization resistance [10]. As a result, the community-level metabolic architecture that governs pathogen resistance in this niche remains uncaptured and therefore poorly defined. Presence or absence of individual taxa alone often does not predict clinical risk. For example, while UPEC is one of the most common etiological agents, together with *Klebsiella pneumoniae*, *Enterococcus faecalis*, and *Proteus mirabilis* [9], increasing evidence links polymicrobial infections to the capacity of uropathogens to persist, interact, and remodel the urinary environment [11]. Notably, organisms such as UPEC and *Enterococcus faecalis* do not behave as isolated invaders: rather, they rapidly reconfigure their metabolic surroundings through cross-feeding, competitive exclusion, and cooperative interactions [9][10][11]. This question becomes particularly crucial in the postmenopausal urinary tract, where communities are typically of low taxonomic complexity yet remain ecologically structured [12]. Whether resistance to recurrent urinary tract infection (rUTI), or its severity, depends on a redundant functional core or on a small set of non-substitutable keystone taxa remains unresolved [11], [15], [16]. In microbial ecology, keystone taxa are defined as community members whose impact on community structure and function is disproportionate to their abundance, such that their removal triggers cascading changes in community composition and metabolic output.

*Winkia anitrata* has emerged as a recurrent but poorly characterized member [17], raising the possibility that specific Actinomycetotal functions may play such a keystone role in the postmenopausal urobiome, despite its reported presence both in commensal urinary communities and in clinical infection settings [18]. To resolve how functions vary, we represent changes in relationships among metabolites across community states as co-variation networks, revealing topological changes that are not apparent from taxonomic composition or metabolite abundance alone.

Addressing this question experimentally requires a readout capable of capturing community-level metabolic organization in its physiological environment and its response to defined compositional perturbations. As argued above, taxonomic profiles and abundance-based metabolomics are insufficient for this purpose [19]: differential metabolite abundance analysis identifies shifts in mean metabolite levels but does not reveal whether the relationships between metabolites have been reorganized. Such relationships can be represented as a metabolite co-variation network, in which nodes are metabolite features and edges denote statistical associations between their abundances across samples [20], [21]. In microbiomes, metabolites that are co-produced, co-consumed, or exchanged tend to vary jointly, forming a structured network of co-variation. When a community transitions between states, the metabolite co-variation network is rewired as edges are gained, lost, or inverted, independently of changes in individual metabolite abundance. We therefore reasoned that metabolite co-variation network rewiring provides a functional readout of community metabolic architecture, reflecting the collective metabolic consequences of microbial perturbations [22].

This concept is operationalized in PhenoRewire, an interpretable network-based framework for metabolite co-variation analysis: it tests each metabolite for association with phenotype or time, reconstructs state-specific Spearman co-variation networks, and quantifies rewiring of co-variation structure between states. As PhenoRewire solely relies on this co-variation structure, it does not require metabolite annotation or prior pathway knowledge and is therefore applicable to partially or fully unannotated feature sets [23]. To enable controlled compositional perturbations, PhenoRewire is coupled with untargeted hydrophilic interaction liquid chromatography–tandem mass spectrometry (HILIC-MS/MS) in two defined-community experiments using iPSC-derived urothelial organoids, a host model that preserves a differentiated urothelial interface capable of sustained urine exposure [24]. This design yields a combined read-out of microbial metabolic dynamics and host metabolic responses during the early phase of colonization.

Here, we leveraged PhenoRewire to interrogate early microbial dynamics in two complementary, defined community models on iPSC-derived urothelial organoids: first, a defined urobiome is challenged with clinically isolated UPEC and *E. faecalis* to model infection; second, a single patient-isolated six-member urobiome is reconstructed and systematically perturbed through single-species dropout. Untargeted HILIC-MS/MS profiling and metabolite co-variation network analysis with PhenoRewire were applied to resolve how uropathogen invasion and community composition reshape metabolic interactions in a physiologically relevant context. Together, these models demonstrate how PhenoRewire can identify emergent community-level metabolic dynamics and reveal the contributions of individual taxa to microbial ecosystem functions, establishing it as a generalizable framework for probing community metabolic dynamics across diverse ecosystems.

## Results

### PhenoRewire identifies distinct metabolic architectures associated with the healthy urobiome

Traditionally, differential abundance testing of individual metabolites is used to assess metabolic differences between biological states. To enable capturing reorganizations in community metabolic architecture across distinct ecological groups associated with healthy and dysbiotic states, we designed PhenoRewire (Fig. 1a).

**Figure 1.**
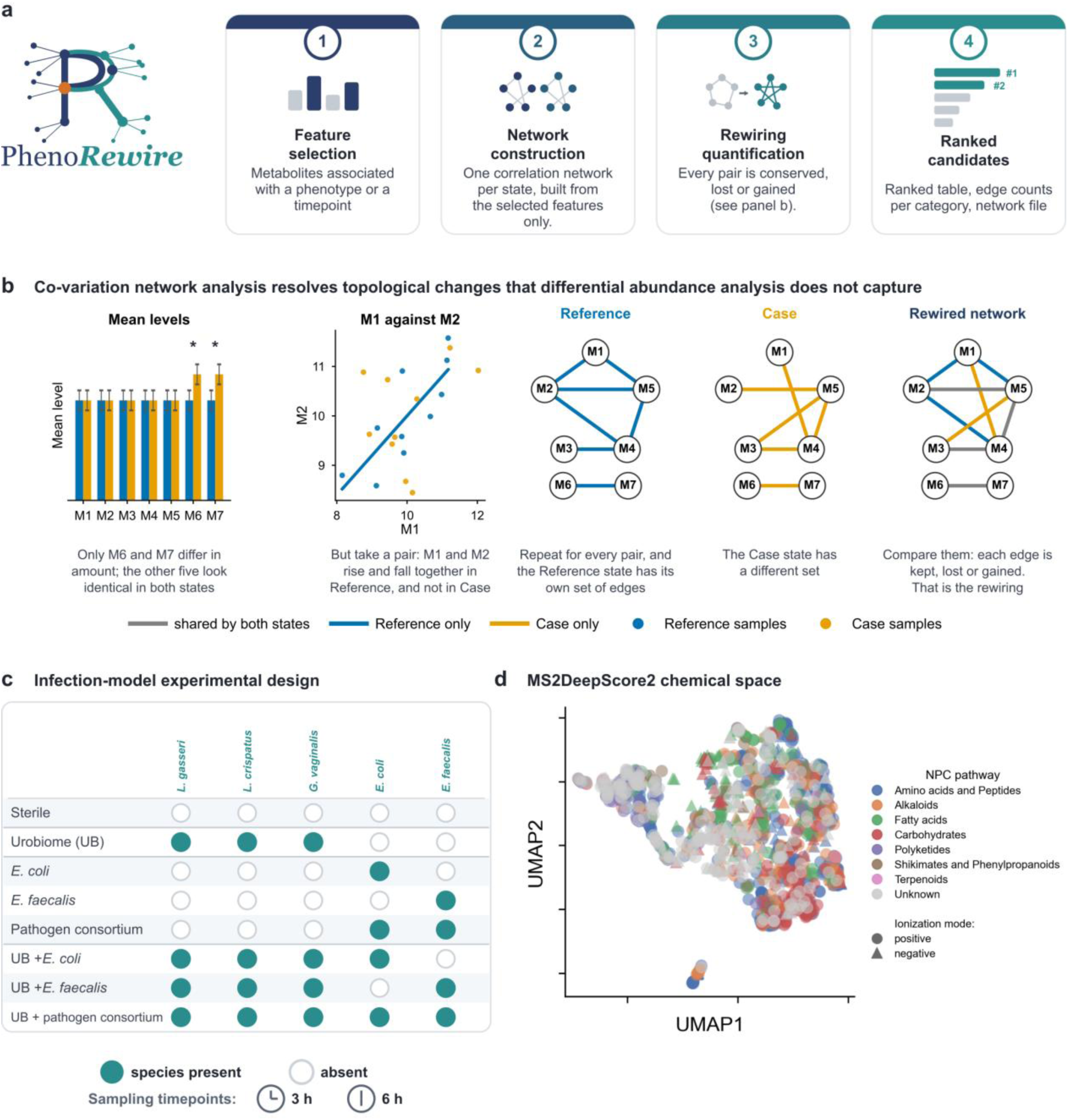
PhenoRewire as a network-based framework for metabolite co-variation analysis, applied to a host–microbiome organoid system. (**a**) Schema of how differential abundance analysis with co-variation network analysis works on a synthetic example. *From left*: mean levels, where only metabolites M6 and M7 differ between the states; M1 plotted against M2, one dot per sample, showing that the pair tracks in the reference but not in the case state; correlation network of each state; rewiring network consisting of the two state networks compared, with each of the nine edges coloured by outcome. (**b**) PhenoRewire workflow: *1.* Feature selection: metabolites are ranked by association with a phenotype (phenotype mode) or a timepoint (temporal mode). *2.* Network construction: one Spearman correlation network is built per state from the selected features, with nodes representing metabolites, sized by abundance. *3.* Rewiring quantification: edges are classified as reference/earlier-state-specific (blue), shared (gray), or comparison/later-state-specific (orange). *4.* Ranked candidates: metabolites are scored by rewiring and exported for network visualization (e.g., in Cytoscape). (**c**) Design of the infection-model experiment: eight defined-community conditions combining the three-member urobiome (*L. gasseri*, *L. crispatus*, *G. vaginalis*) with the uropathogens *E. coli* and *E. faecalis*, alone or in combination; filled circles indicate species present, open circles absent. Apical medium was sampled at 3 h and 6 h. (**d**) MS2DeepScore2-based UMAP projection of annotated and CANOPUS-predicted metabolite features of the infection model, colored by NPC compound class pathway and shaped by ionization mode. Co-localization of metabolites within the same macro-class, together with overlapping distributions across ionization modes, supports the chemical coherence of the classification and complementary polarity coverage.

PhenoRewire evaluates whether the co-variation structure among metabolites is altered, in one of two modes. In phenotype mode, two biological states are compared at a fixed timepoint to test whether compositional differences reorganize the co-variation architecture. Alternatively, in temporal mode, the framework compares two timepoints within the same condition, examining how a community’s metabolic architecture evolves. This dual-mode approach enables the systematic deconvolution of context-dependent metabolic dependencies, distinguishing between transient, community-intrinsic temporal shifts and persistent perturbation-induced structural reconfigurations (Fig. 1b). PhenoRewire operates directly on standard untargeted metabolomics outputs, metabolite intensity and sample metadata tables, common to any LC-MS/MS pipeline [25]. It remains compatible with unannotated metabolites and requires no pathway mapping.

For each comparison, features are ranked by how strongly their co-variation neighborhood differs between the two states, and a correlation network is reconstructed per state from the selected features (Fig. 1b). As this ranking-based selection drives feature inclusion, rather than a fixed cutoff, even comparisons with limited statistical power yield interpretable networks when a genuine signal is present. In the next step, comparison of the two networks enables the classification of every connection as shared, specific to one state, or sign-reversing (Fig. 1a). In the resulting rewiring visualization, node size encodes each metabolite’s rewiring score, while node and edge color denote the state association. Network-level modularity helps distinguish sparse, coordinated architectures from dense, indiscriminate ones and is flagged as unreliable if it falls below a fixed threshold (see Methods). Independently of the network, each run also generates a ranked metabolite table for every comparison, containing group means, fold-change, and rewiring scores for all selected features, usable as a standalone result alongside GraphML networks and Markdown/JSON summary reports (full specifications in Methods and Supplementary Methods SM1–SM4).

To test PhenoRewire, we applied the pipeline to a host-microbiome system using iPSC-derived urothelial organoids [24], [26], [27] across defined-community experiments designed to probe perturbations in composition and over time. First, we questioned how interactions between invading uropathogens *E. coli* and *E. faecalis* and a defined urobiome reshape the metabolic architecture of the community during early colonization (Fig. 1c). This ‘healthy’ urobiome consisted of *L. gasseri* and *L. crispatus,* commonly associated with reduced rUTI risk [12], together with *G. vaginalis,* a taxon linked instead to UTI recurrence [28]. Samples were collected at 3 h and 6 h, timepoints capturing initial niche colonization dynamics [29], and metabolites were annotated via a six-tool integrative pipeline at MSI confidence levels 1–4, yielding 176 and 287 features confidently annotated at MSI levels 1–2 across both ionization modes in the infection and strain-dropout experiments, respectively.

Subsequently, we used CANOPUS compound class predictions to extend chemical characterization, enabling characterization of 70.3% of features. This coverage, spanning amino acids, peptides, and polyketides, supported biochemical interpretation for the majority of the metabolome, including unannotated features. The coherence of this classification was validated by MS2DeepScore2-based UMAP projections [30], where metabolites within the same macromolecular class showed preferential co-localization, and the positive and negative ion mode features that overlapped in chemical space confirmed complementary polarity coverage. (Fig. 1d).

We first asked which community members of the infection model drove metabolic differentiation from the sterile organoid control, and whether this changed between 3 and 6 h. Global multivariate analysis revealed a pronounced temporal inversion in the primary driver of metabolic differentiation from the sterile organoid control (Supplementary Table S1). At 3 h, urobiome-containing conditions were the only ones significantly distinct from the sterile organoid control (PERMANOVA positive mode: pseudo-*F* = 3.65, *P* = 0.003), while *E. coli*, *E. faecalis*, and the pathogen consortium remained indistinguishable from the sterile organoid control. By 6 h, this ordering had fully reversed: *E. coli* emerged as the primary driver of metabolic divergence (PERMANOVA negative mode: pseudo-*F* = 9.84, *P* = 0.002), while the urobiome was no longer separable from the sterile control. The pathogen consortium followed the same delayed trajectory as *E. coli* monoculture, reaching significance only at 6 h, consistent with *E. coli*, rather than *E. faecalis*, as the driver of the late-phase metabolic response. Copresence configurations reinforced this interpretation: all three combinations containing the urobiome (urobiome with *E. coli*, *E. faecalis*, or both) separated from sterile control conditions at 3 h, whereas *E. coli* and *E. faecalis* monocultures did not. Early differentiation therefore tracked the presence of the urobiome rather than the presence or identity of a pathogen. PLS-DA score plots supported this temporal trend, showing the same early separation of urobiome-containing conditions and the later divergence driven by *E. coli* (Fig. 2a, b), and identified the metabolic features underlying this inversion. At 3 h (LV1 = 12.4%, LV2 = 7.0% of total X variance), urobiome-containing communities occupied a distinct region of score space while *E. coli*, *E. faecalis*, and sterile conditions clustered together (Fig. 2a, Supplementary Fig. S2). This early divergence was driven by a coherent set of top-ranked metabolites based on variable importance in projection (VIP) loadings, whereby higher VIP scores indicate a stronger contribution to the class separation (Supplementary Table S2).

**Figure 2.**
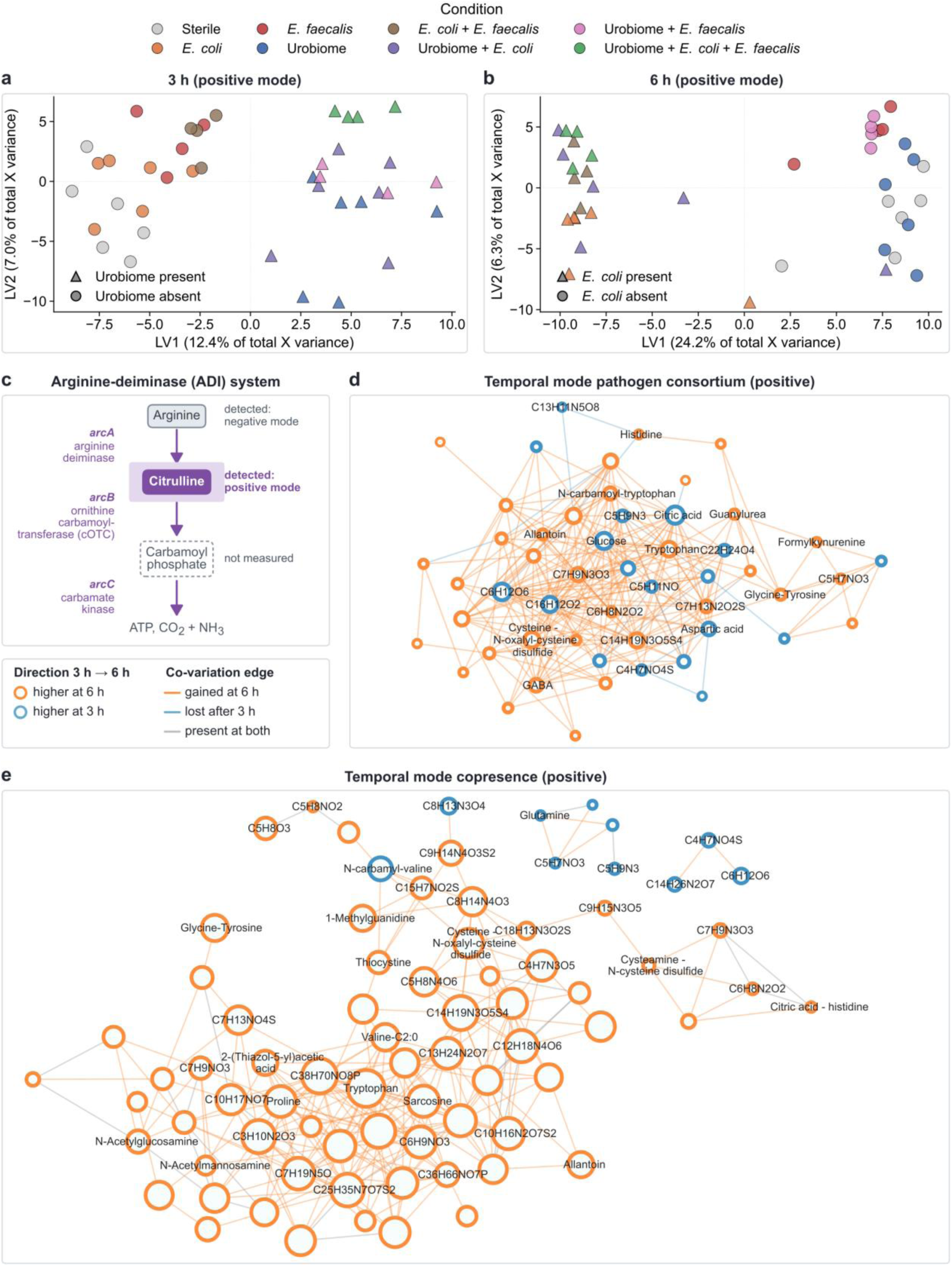
The urobiome, uropathogens, and their copresence generate distinct temporal metabolic architectures in the iPSC urothelial organoid infection model. (**a,b**) PLS-DA score plots (positive ion mode) at 3 h (a) and 6 h (b), with samples colored by experimental condition. Axis labels report each latent variable’s (LV) score variance as a percentage of total variance in the preprocessed feature matrix. At 3 h, urobiome-containing communities (triangles) separate from all other conditions along LV1; by 6 h, *E. coli*-containing conditions (triangles) drive separation along LV1, while urobiome and sterile samples converge. (**c**) The arginine-deiminase (ADI) pathway: citrulline was differentially abundant and rewired in 6 of 21 PhenoRewire comparisons (4 at q < 0.05; strongest q = 0.008, pathogen consortium vs. copresence, 3 h), while arginine was detected only in negative mode. (**d**) Temporal co-variation network for pathogen consortium condition showing the transition from a structure anchored by citric acid, glucose, and aspartic acid, defining a sparse carbon/TCA-linked network to a substantially denser network enriched in amino acid and sulfur/thiol-containing species including GABA, histidine, tryptophan, cysteine-N-oxalyl-cysteine disulfide, and thiocystine, consistent with a transition from oxidative carbon metabolism to redox-adaptive amino acid co-regulation. (**e**) Temporal co-variation for copresence condition, coloured as in d, showing a more coordinated carbamoyl/nitrogen signature at 3 h, and forming a dense co-variation cluster dominated by amino acid, nitrogen, and sulfur/thiol-linked species, showing an 8-fold network expansion.

Citrulline (VIP = 3.30;), N-carbamoyl-proline (3.27), ureidosuccinic acid (3.22), nicotinic acid (3.21), N-carbamoyl glutamic acid (2.81), and gamma-aminobutyric acid (GABA) (2.69) all showed high VIP scores and converge on a coherent carbamoyl-phosphate metabolic signature. We next asked whether the urobiome’s own co-variation architecture changed over the same interval. Phenorewire analysis indicated that the urobiome network density increased only modestly between 3 h and 6 h, and the dominant co-variation metabolites remained largely conserved, with limited expansion toward histidine and kynurenine at 6 h (Supplementary Figure S3). This suggests the healthy urobiome undergoes early community-level adaptation that reshapes nitrogen and pH-related metabolism before the pathogen-specific metabolism emerges. Specifically, the arginine deiminase (ADI) pathway appears to be activated (Fig. 2c), whose products citrulline, ornithine, and ammonia are known to modulate uropathogen fitness [31], [32]. This interpretation is further supported by network-level rewiring. Citrulline and homocitrulline were differentially abundant in 6 of 21 PhenoRewire comparisons and specifically in urobiome, pathogen, and copresence conditions (best q = 0.008), while N-carbamoyl-proline and ureidosuccinic acid were rewired consistently regardless of ionization mode or comparison type, indicating systematic perturbation of carbamoyl phosphate–linked metabolism. Together, these results suggest that the urobiome establishes a stable carbamoyl-nitrogen metabolic signature and that this bacterial carbamoyl-phosphate flux contributes to reshaping the host N-carbamoyl metabolite pool.

### Uropathogens drive progressive metabolic rewiring toward sulfur and amino acid co-variation that is further amplified by E. faecalis

As the presence of pathogens (*E. coli* or *E. faecalis* monocultures, and the pathogen consortium) drove the most pronounced temporal rewiring, we next investigated the pathogen-induced metabolic rewiring in more detail. In the absence of the healthy urobiome, the co-variation structure at 3 hours was anchored by citric acid, glucose, and aspartic acid, defining a carbon/TCA-linked network consistent with active oxidative metabolism during early community establishment. By 6 h, this architecture underwent an extensive transition toward a denser network dominated by GABA, histidine, tryptophan, and thiol-containing species including cysteine-N-oxalyl-cysteine disulfide and thiocystine (Fig. 2d; Supplementary Fig. S4). This shift toward sulfur/amino acid co-regulation likely reflects a niche adaptation to the oxidative stress in the urothelial environment: uropathogenic *E. coli* encounters reactive oxygen and chlorine species during urothelium colonisation, and cysteine thiol oxidation serves as primary target of this adaptive response [33].

To identify the microbial driver of this rewiring, *E. coli* and *E. faecalis* were analyzed as monocultures (Supplementary Fig. S5). *E. coli* monoculture reproduced the directional co-variation architecture of the consortium, establishing *E. coli* as the primary architect of the pathogen temporal signal. Yet, the consortium showed substantially amplified rewiring compared to the *E. coli* monoculture, showing an approximately a 6-fold increase in edge count in the presence of *E. faecalis* (Supplementary Table S3). In contrast, *E. faecalis* monoculture yielded no differentially abundant features at either timepoint or ionization mode, and its standalone co-variation architecture was indistinguishable from the host-only background.

Thus, *E. faecalis* contributes to infection not through an independent metabolic footprint, but through network-level facilitation that manifests exclusively in the consortium context. Together, these data suggest that while *E. coli* initiates the metabolic rewiring, the full extent of the reorganization is a synergistic emergent property that cannot be reduced to the independent activity of either organism alone.

To understand the influence of urobiome–pathogen copresence on the metabolic rewiring of the community as a whole, PhenoRewire analyses was run in temporal mode on the combined healthy and pathogenic community. We observed that despite simultaneous inoculation of all conditions from freshly prepared inoculum, the urobiome-pathogen copresence co-variation network was already substantially more densely connected than the pathogen network at 3 h (Supplementary Table S3). Between 3 h and 6 h, the urobiome–pathogen copresence condition showed the strongest temporal rewiring of all conditions: network density expanded, forming a dense amino acid/nitrogen/sulfur cluster comprising histidine, tryptophan, allantoin, N-acetylglucosamine, cysteine-N-oxalyl-cysteine disulfide, and 1-methylguanidine (Fig. 2e). This late co-variation architecture is ∼8-fold greater in edge count than the pathogen late network (1,781 vs 227 positive-mode edges at the applied correlation threshold) and is not recapitulated by either individual community, establishing copresence as an emergent metabolic state.

To quantify what co-variation analysis adds over abundance testing in this experiment, we compared the two readouts directly. Classical differential-abundance analysis identified 259 in positive and 75 in negative mode features, while PhenoRewire analysis additionally identified 227 in positive and 66 in negative ion mode metabolites rewired across all condition-pair comparisons, identifying 3,437 co-expression edges (2,632 gained, 804 lost, 1 sign-switched) that are not evident from standard differential abundance testing showing that metabolite abundance and co-variation capture complementary information (Supplementary Table S8). Overall, PhenoRewire established that our defined three-member microbial community [34] (*L. crispatus*, *L. gasseri*, and *G. vaginalis*) generates a characteristic co-variation architecture that invading uropathogens reorganize but do not reproduce upon *E. coli* invasion, particularly in the presence of *E. faecalis*.

### Compositional perturbation resolves the Winkia anitrata dropout as the sole outlier, whose removal causes near-total collapse of community co-variation topology in a six-species urobiome community

To better understand how different members of a more complex, patient-isolated community shape the characteristic co-metabolic architecture, we performed a strain-dropout experiment. To this end, we assembled a six-member community reconstructed from the urinary microbiome of a postmenopausal donor that consisted of *Winkia anitrata*, 2 *Enterococcus faecalis* strains, *Streptococcus vaginalis*, *Staphylococcus pasteuri*, and *Staphylococcus epidermidis*. Next, we applied systematic single-species dropout across all members and assessed, for each dropout, how removing a single species altered community metabolic profiles and co-variation network structure, enabling the identification of members whose absence disproportionately disrupts the community (Fig. 3a). To enable a direct comparison of co-variation architecture between this composition-driven perturbation and the infection-driven perturbations described above, the analytical platform, timepoints, and analytical pipeline were kept identical. Structural annotation combined with CANOPUS compound class predictions yielded 60.3% classification, with UMAP projection confirming preferential co-localization of metabolites within the same macromolecular class (Fig. 3b). Classical differential-abundance analysis identified 247 significant features in negative (q < 0.05), and none in positive ion mode, whereas PhenoRewire analysis identified 134 in negative and 243 in positive ion mode metabolites rewired within all the strain-dropout comparisons, detecting a total of 3,282 rewired feature-comparison instances, an approximately 13-fold increase in detection range (Supplementary Table S8).

**Figure 3.**
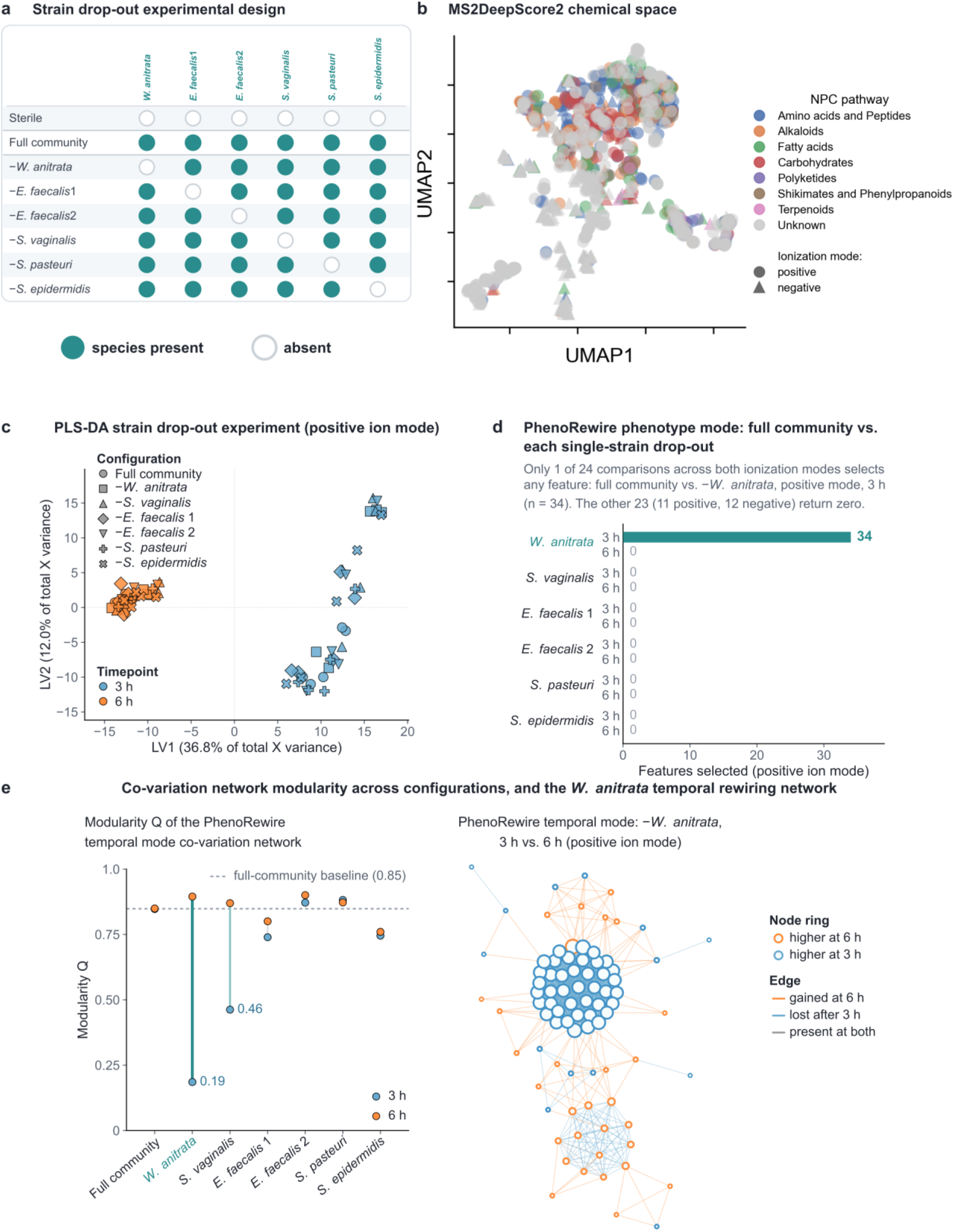
*W. anitrata* is a keystone species that uniquely reorganizes the six-member urobiome’s temporal co-variation architecture. (**a**) Composition of the strain-dropout experiment: a full six-member consortium (*W. anitrata*, *E. faecalis* strain 1, *E. faecalis* strain 2, *S. vaginalis*, *S. pasteuri*, *S. epidermidis*) with each member sequentially dropped, across 8 conditions. (**b**) MS2DeepScore-based UMAP projections of annotated and CANOPUS-predicted metabolite features for the strain-dropout experiment, colored by NPC compound class pathway and shaped by ionization mode. Co-localization of metabolites within the same macro-class, together with overlapping distributions across ionization modes, supports the chemical coherence of the classification and complementary polarity coverage. **(c**) PLS-DA of the strain drop-out experiment in positive ionization mode, both timepoints (LV1 = 36.8%, LV2 = 12.0% of total X variance). Samples are colored by timepoint and shaped by configuration. Metabolic profiles separate primarily by timepoint rather than by community configuration, and converge by 6 h regardless of which species is absent. **(d**) PhenoRewire phenotype-mode comparisons of each single-species dropout vs. the full community; of 24 total comparisons across both ionization modes, only −*W. anitrata* vs. full community at 3 h returns selected features (n = 34) and a rewired network; all the remaining returned zero. **(e**) *Left*, modularity (Q) of the temporal co-variation network for each configuration, in positive ionization mode; the full community and four single-species dropouts remain close to the full-community modularity baseline at both timepoints; -*S. vaginalis* shows a partial, statistically non-significant reduction at 3 h (Q = 0.46); *−W. anitrata* collapses toward near-random co-variation at 3 h (Q = 0.19) before reorganizing by 6 h (Q = 0.90). *Right,* the −*W. anitrata* temporal rewiring network underlying this collapse: nearly all 112 temporally selected features form a single densely interconnected cluster visually consistent with Q = 0.19, while the 6 h-specific connections (orange) begin to differentiate the network into more distinct groups.

We first assessed whether community composition or sampling time dominated overall metabolic variance in this experiment. Global multivariate analysis indicated that timepoint was the dominant axis of metabolic variance across all distance metrics and ionization modes (Supplementary Table S4).

At 3 h, community composition was a statistically significant predictor of metabolic profiles; by 6 h the composition effect disappeared entirely in positive mode (all metrics *P* > 0.15), consistent with metabolic convergence (Fig. 3c), while a residual compositional signal persisted in negative mode (Bray-Curtis: pseudo-*F* = 1.99, *P* = 0.012), indicating that anionic metabolites retain strain-specific community information at the later timepoint (Supplementary Fig. S6, Supplementary Table S5).

We then tested which single-species removals altered community co-variation relative to the full community. Removing *W. anitrata* was the only single-species perturbation that reorganized community metabolic co-variation, as evidenced by the phenotype mode comparisons, whereby only the full community versus *W. anitrata*-absent returned phenotype-associated features (n = 34; FDR = 0.20, pre-specified ceiling) (Fig. 3e). For the other species, no metabolites were identified that discriminated the dropout from the full community, indicating that their co-variation landscapes are statistically indistinguishable from the full community. The metabolic equivalence of the two *E. faecalis* dropouts is also in line with comparative genomics of the community isolates, which shows the two *E. faecalis* strains to be near-identical (digital DNA-DNA hybridization 99.8%; COG-content Jaccard similarity 0.991; only 13 accessory orthologous groups distinguishing them).

In line with the phenotype comparisons, PhenoRewire temporal analysis in positive ion mode showed that the co-variation network modularity (Q) remained stable in the full community between 3 h and 6 h (Q ∼ 0.85), whereas the *W. anitrata*-absent network collapsed toward a near-random topology at 3 h before reorganizing by 6 h (Q from 0.19 to 0.90; Fig. 3e). In accordance, when looking at the extent of connectivity, the *W. anitrata*-absent community lost more connectivity than any other dropout configuration (from 958 to 144 edges, -85%), a result reproduced in negative ion mode (Supplementary Table S6). In contrast, other members yielded heterogeneous, often contradictory responses: the *S. pasteuri*-absent network gained connectivity (from 76 to 170 edges, +124%), whereas in positive mode *S. vaginalis* removal produced a partial and statistically non-significant reduction in modularity (Q = 0.46 at 3 h), consistent with a secondary role in the community. Co-variation architecture was also preserved across all five single-dropout conditions involving strains (Supplementary Table S6). This indicates the absence of a generic dropout-induced network collapse effect and instead identifies *W. anitrata* as the keystone member governing temporal network dynamics. Indeed, the network consequences of *W. anitrata* depletion were significant. While the full community maintained a sparse, modular co-variation network (34 selected features, 23 edges, Q = 0.707), *W. anitrata* removal induced a radical topological shift: the resulting network comprised 258 edges with Q = 0.038, approaching the theoretical minimum for random co-variation (below the Q ≥ 0.3 threshold for module-level inference; see Methods). This global rewiring proportion was 96.7%, with 263 of 272 union edges being state-specific and only 9 shared between conditions. In contrast, all five other single-species dropout communities retained a network topology statistically indistinguishable from the full community, further underscoring *W. anitrata* as the keystone species.

To understand the metabolic phenotype induced by *W. anitrata*, we investigated the 34 features selectively enriched in the *W. anitrata*-absent community in more detail.

These metabolites span distinct functional classes and include the aromatic amino acids tryptophan and tyrosine, the amino sugars 2-acetamido-2-deoxy-β-D-glucosylamine and galactosamine, the porphyrin precursor 5-aminolevulinic acid, and metabolites of central carbon and nitrogen metabolism including citric acid, methylguanidine, and pyridoxamine. The most interpretable axis observed in positive ionization mode involved aromatic amino acid catabolism. We observed a co-enrichment of tryptophan and tyrosine, which is consistent with impaired aromatic catabolism in the absence of *W. anitrata*. The parallel enrichment of N-carbamoyl-tryptophan suggests that part of the accumulating tryptophan pool is diverted towards carbamoylation. Pyridoxamine, as the dephosphorylated form of pyridoxamine 5′-phosphate, the direct product of pyridoxal 5′-phosphate-dependent transamination reactions, could point to reduced aminotransferase-linked nitrogen cycling. Citric acid and methylguanidine indicate secondary TCA remodeling and altered arginine-creatinine catabolism.

N-acetyl-D-mannosamine, a sialic acid precursor and glycan recycling intermediate, showed the strongest temporal signal of all annotated negative-mode features, accumulating at 3 h and disappearing entirely by 6 h. This profile is consistent with impaired processing of urothelial glycocalyx-derived sugars in the absence of *W. anitrata* [35]. Allantoin showed the same temporal arc, consistent with a transient oxidative challenge progressively attenuated by remaining members. Collectively, these patterns implicate *W. anitrata* in aromatic amino acid catabolism, oxidative-purine turnover, and host-interface glycan processing. Critically, the *W. anitrata* dropout produced no differentially abundant metabolites relative to the full community, yet it exhibited the highest per-strain rewired metabolites count (90), illustrating that its influence operates through co-variation topology rather than mean metabolite levels.

### The five Bacillota are metabolically redundant, and a late-phase composition-independent program persists

To investigate to which extent the five Bacillota species, together with *W. anitrata*, were metabolically redundant in the six-species urobiome community, we performed leave-one-out (LOO) attribution across the whole experiment, quantifying each strain’s contribution to community-wide co-variation. To this end, each rewired metabolite was assigned to the community member whose removal most reduced its rewiring score relative to the full community. Of the 372 metabolites tested, 196 (53%) were attributable to the removal of a single strain, while the remaining 176 showed no single-strain-dependent loss. Among attributed metabolites, *W. anitrata* (90) and *S. vaginalis* (76) jointly accounted for 85% of single-member attributions, while the four remaining Bacillota strains (*E. faecalis* strain 1 (11), *S. pasteuri* (9), *E. faecalis* strain 2 (8), and S*. epidermidis* (2)) contributed comparatively few. Thus, although only *W. anitrata* removal reorganizes the phenotype-associated network architecture itself, per-feature attribution identifies both *W. anitrata* and *S. vaginalis* as disproportionate contributors to community-wide co-variation, establishing *S. vaginalis* as a genuine secondary contributor rather than an interchangeable member. This distributed contribution followed a distinct temporal trajectory from that of *W. anitrata*. Classical differential abundance testing detected no significant *S. vaginalis*-associated features at 3 h but 36 features at 6 h (negative mode, q < 0.05), spanning aromatic/indole derivatives (indole-3-acetic acid, 3-acetylindole), ADI-pathway/polyamine intermediates (homocitrulline, spermine, arginine), and TCA-linked species (citric acid), several of the same functional classes altered in the *W. anitrata*-absent community.

Unlike *W. anitrata*, whose absence reorganized network topology from the earliest timepoint rather than changed metabolite abundances, S*. vaginalis*’s broader, later-onset metabolic footprint left community modularity essentially unaffected.

We last asked whether any part of the metabolic program persisted independently of community composition. Finally, a late-phase metabolic program emerged at 6 h across community compositions regardless of which species was absent. For example, the sulfonated feature C_8_H_6_O_6_S (m/z 248.02), increased from 3 h to 6 h in the full community and in all six single-strain dropouts, including the *W. anitrata*-absent condition. Two further features followed the same 6 h direction in most but not all compositions: C_12_H_17_N_5_O_9_S_2_ (m/z 478.01) increased in the full community and five of six dropouts, being absent from the selected set only in the *E. faecalis* strain 1-dropout, while C_7_H_19_N_3_O_4_S (m/z 206.09) likewise increased across the full community and five dropouts but was absent specifically from the *W. anitrata* one. The robustness of this late program to single-species loss indicates that it is encoded redundantly across the community, in contrast to the strain-specific early co-variation architecture. These observations establish a structural dissociation in community metabolic organization: *W. anitrata* disproportionately shapes early-state co-variation topology, a role no other single member recapitulates, whereas the late-phase metabolic trajectory is a distributed community property robust to single-species loss.

## Discussion

The urinary microbiome regulates urothelial homeostasis through community-level metabolic interactions, yet most studies remain taxonomy- and genome-centred, inferring function from composition or encoded potential rather than capturing the metabolome as an organized regulatory system. This distinction matters because communities undergo state transitions - pathogen invasion, keystone species loss, temporal maturation - and may reorganize co-variation architecture without proportionally altering individual metabolite abundances. Co-variation network analysis complements the abundance-based readouts that dominate microbiome research. By testing whether metabolite relationships are reorganized rather than whether mean metabolic levels shift, it recovers state transitions that differential abundance leaves invisible. This distinction is biologically relevant: in our infection model at 6h, the 3 member urobiome condition showed higher levels of hypoxanthine, tyrosine, citric acid, and glucose than the pathogen condition, yet it was the pathogen network (*E. coli*, *E. faecalis*, and co-colture) that was more extensively reorganized. In this context, PhenoRewire provides a state-aware view of metabolic organization that is complementary to genome-based or taxonomical approaches.

PhenoRewire has several characteristics that make it particularly suited for investigating microbiomes as dynamic, adaptive systems rather than static compositions. First, PhenoRewire does not require pathway annotations, organism-specific databases, or prior knowledge of metabolite identity: co-variation structure is detectable in partially or wholly unannotated feature sets, making the framework applicable to any multi-condition untargeted metabolomics dataset. The framework generates ranked, testable hypotheses rather than binary significance calls, and is designed to be reused across experimental layouts through a YAML configuration file without modifying the underlying code. Thus, PhenoRewire can be integrated as an orthogonal, interpretable layer against which to validate observations drawn from other data types in multi-omics experimental set-ups. At the same time, because PhenoRewire’s inference depends exclusively on metabolomic co-variation, the biological interpretability of its output is bounded by the annotation quality of the input data: a network built entirely on unannotated features can flag rewiring but cannot explain it, risking interpretations built on structurally real yet biologically spurious correlations. Here, the use of our six-tool annotation pipeline preceded co-variation network analysis, which allowed us to establish chemical coverage through direct library matches, analogue search, or compound-class predictions, and anchored prioritized network nodes to plausible biochemistry.

Existing computational approaches generally converge on three distinct paradigms. First, differential-correlation-network methods developed for genome-scale co-expression data, including DGCA, discordant, and DiffCoEx [36], [37], [38] enable the detection of condition-dependent changes in a pairwise or module-level correlation structure. However, these methods were designed and validated exclusively on gene identifiers and have not been benchmarked on untargeted metabolomics datasets, where metabolite identity is frequently unknown. Second, within metabolomics specifically, several methods facilitate feature comparison across biological states, but all have substantial drawbacks.

DNEA supports unannotated, untargeted feature sets, but its regularization is calibrated for 50–500 samples per group and it is restricted to two-group comparisons. BioNetStat supports comparisons across more than two biological states, but requires features to be pre-assigned to defined variable sets (e.g., pathways or ontology terms), precluding its use on unannotated, untargeted feature sets [39]. DiffCorr tests pairwise correlation differences rather than generating a per-metabolite ranking suitable for hypothesis generation [40]. MetVAE estimates sparse metabolite correlation matrices from untargeted data, adjusting for compositionality, missingness and confounding, and has been applied across disease states, but estimates one network per dataset without edge-level classification or per-feature ranking [41]. Third, network-based approaches are used to analyze untargeted metabolomics data, including molecular networking tools such as GNPS2 [42]. However, these tools were designed to support annotation and dereplication by connecting features through MS/MS spectral similarity, a fundamentally different objective that does not test whether co-variation structure differs between biological states. PhenoRewire is therefore the first framework to combine annotation-agnostic feature handling, flexible multi-state comparison, and per-feature rewiring ranking in a single design.

To demonstrate the power of PhenoRewire, we here applied the tool to show how metabolic co-variation network analysis captures early metabolic transitions in both a reconstructed urobiome and an infection model. In our iPSC-derived urothelial organoid infection model, we observed that healthy urobiome species induced a temporally stable extracellular co-variation of citrulline, ureidosuccinic acid, N-carbamoyl-proline, and N-carbamoyl glutaric acid, suggesting the urobiome species are actively establishing a metabolically distinct niche during early colonization. These metabolites are connected through the arginine deiminase (ADI) pathway, a conserved energy-generating route in Lactobacilli in which arginine is converted to citrulline and ammonia, with citrulline and ornithine actively exported via dedicated antiporters [43], [44], [45] and ureidosuccinic acid, N-carbamoyl-proline, N-carbamoyl glutaric acid serving as downstream carbamoyl phosphate overflow products. Citrulline and ureidosuccinic acid are themselves key intermediates of the arginine/citrulline cycle and pyrimidine biosynthesis, both downstream of carbamoyl phosphate. Both *L. gasseri* and *L. crispatus*, two of the three defined urobiome members in our model, are established urogenital colonizers in lineages with genomically conserved ADI pathway genes [46], [47]. Extracellular citrulline accumulation is a well-documented marker of ADI activity [44], and the co-variation of carbamoyl phosphate overflow products is consistent with sustained ADI flux, with carbamoyl phosphate partitioning between phosphoryl transfer to ADP by carbamate kinase, de novo pyrimidine biosynthesis via ureidosuccinic acid [48], [49] and non-enzymatic amino acid carbamoylation. Interestingly, L-ornithine, the terminal product of the ADI pathway, can also be exported by *E. faecalis* and has been shown to serve as metabolic cue that facilitates *E. coli* siderophore biosynthesis and iron acquisition during polymicrobial infection [50], while the ammonia released by the same pathway can raise local pH and restrict urothelial colonization [31], [32]. Ornithine was not directly detected in our dataset, so this cross-feeding remains a plausible interpretation supported by the observed metabolite pattern.

Thus, the urobiome ADI pathway may serve dual roles: in Lactobacillus, arginine catabolism via ADI yields ATP, supporting their growth and niche colonization, while the concomitant export of ornithine may provide a metabolic cue that promotes iron acquisition in co-colonizing Enterobacteriaceae.

In the same model in the absence of a healthy urobiome, the uropathogens *E. coli* and *E. faecalis* displayed a transition from carbon/TCA-centred co-variation (citric acid, glucose, monosaccharides at 3 h) to sulfur/amino acid-enriched co-variation (thiocystine, cysteine-N-oxalyl-cysteine disulfide, histidine, tryptophan at 6 h). This is consistent with uropathogens adapting to host-associated stress during infection, where early aerobic central carbon metabolism transitions to thiol-based redox homeostasis [50]. The *E. coli* monoculture reproduced this directional architecture, confirming *E. coli* as the primary mediator. The observed network-level amplification in the presence of *E. faecalis* is consistent with ecological facilitation, with previous studies showing that *E. faecalis* can promote *E. coli* persistence through suppression of bladder innate immune responses during co-infection [51], and enhancement of biofilm formation and antibiotic recalcitrance in the catheterized urinary tract [52].

Culturing the defined urobiome with one or both uropathogens triggered an extensive metabolic reorganization within the infection model. By 3 h, the copresence network already carried the urobiome’s carbamoyl-nitrogen signature regardless of pathogen identity, suggesting the urobiome establishes a constitutive metabolic program early in community assembly. However, this pattern was inverted by 6 h, as copresence networks exhibited roughly 8-fold higher edge count than the pathogen-alone networks and an increase in inter-timepoint connectivity. This hyper-connectivity supports two non-exclusive, yet mechanistically distinct, hypotheses. First, the network rewiring may facilitate pathogen activity. Urobiome-derived ornithine via the ADI pathway may for example cross-feed and potentiate *E. coli* iron-acquisition and stress-response metabolism. Alternatively, the network amplification could reflect an intensified metabolic contest, in which the host organoid and urobiome mount a multifaceted response to pathogen invasion rather than a unilateral facilitation of pathogen activity. Whereas our data unequivocally establish that copresence heightens metabolic reorganization, determining whether this intensification represents synergy or competition is beyond the capabilities of purely co-variation network analysis. Resolving these alternatives would require integrating PhenoRewire outputs with longitudinal pathogen burden and host-response readouts. Moreover, also the iPSC-derived urothelial organoids present in this model inherently participate in local arginine metabolism, adding a layer of host-derived metabolic complexity to this ecological niche. Upon air-liquid interface differentiation, these cultures form a polarized epithelial barrier and strongly upregulate NOS1 and NOS3, which synthesize NO from L-arginine with L-citrulline as co-product. Concomitantly, ASS1 is upregulated, the rate-limiting enzyme that regenerates arginine from citrulline by condensing it with aspartate to form argininosuccinate, thereby sustaining NO synthesis [53], (Bindas et al., manuscript in preparation).

Another interesting metabolic axis across both our infection model and the dropout model involves allantoin, a purine catabolism product of non-enzymatic ROS-mediated oxidation of uric acid. Allantoin provides the clearest cross-community signal in this dataset: it recurs as a component of the late pathogen co-variation network in the infection model and as an early-accumulating feature in *W. anitrata*-absent communities in the dropout model. Three non-mutually-exclusive mechanisms could account for this: urothelial oxidative stress via UPEC-mediated NRF2 suppression driving uric acid oxidation [54]; *E. coli* nitrogen scavenging via the allantoin regulon [55]; and a possible *W. anitrata*-dependent catabolism of this substrate in the intact community. Resolving these hypotheses into relative contributions requires isotope-traced supplementation and ROS quantification. The concomitant accumulation of tryptophan and tyrosine in *W. anitrata*-absent communities is consistent with reduced aromatic amino-acid turnover, although direct biochemical evidence for *W. anitrata*-dependent aromatic catabolism remains to be established. For the host, the accumulation of tryptophan in the pathogen late-phase network is biologically significant to urothelial immunity, given that indole derivatives and tryptophan catabolites are established suppressors of NF-κB signalling and promoters of immune tolerance in epithelial cells [22], [56]. In parallel, the enrichment of allantoin may reflect a community state characterized by altered nitrogen/purine handling and increased oxidative stress, pointing to a broader destabilization of community metabolism rather than a single-pathway effect. The early hyperconnected, low-modularity network of *W. anitrata*-absent communities compared to the stable modular full community indicates that loss of this phylogenetically distinct Actinomycetota member of an otherwise Bacillota-dominated consortium destabilizes co-regulation specificity as its primary network consequence, suggesting a community organizer function that is mechanistically distinct from, and precedes, its substrate consumption roles [57], [58], [59]. This distinction also clarifies what keystone means functionally in this model, in line with the classical ecological definition of a keystone species: impact on community structure disproportionate to abundance, such that removal triggers cascading reorganization [16]. The number of metabolites attributed to a strain therefore reflects the breadth of its metabolic footprint, but not its structural centrality to community organization. *S. vaginalis*, whose metabolic reach overlaps substantially with that of *W. anitrata*, is a broad but structurally dispensable contributor: only *W. anitrata* removal triggers the cascading reorganization that defines a structural keystone.

Our study has several limitations. First, the urothelial iPSC–urobiome model used here is, by design, a simplification of the *in vivo* urinary tract. Yet within the current state of the art it represents a meaningful advance: more controlled and reproducible than clinical urine sampling, more physiologically relevant than monoculture or broth, and uniquely able to decompose community-, strain-, and host-level contributions through defined dropout and copresence designs. Barrier cultures from the same iPSC-derived organoid lineage strongly resist 40-kDa fluorescent dextran transport and mount IL-6 responses upon 24-hour UPEC challenge (Bindas et al., manuscript in preparation). Moreover, to our knowledge, this is the first study to co-culture a defined, single patient-isolated urobiome community with a host urothelial model *in vitro*. Existing organoid and bladder-on-a-chip systems have modeled infection with single uropathogens [24], [60], while defined urobiome consortia have been studied without a host compartment [61].

Although combining a urobiome community with a host urothelial model represents a substantial advancement, the model lacks the immune compartment, mechanical flux of micturition, and mucosal heterogeneity. Second, untargeted metabolomics does not provide absolute quantification, and supernatant metabolomics cannot fully discriminate host-derived from microbial metabolites. All inferences are associative and do not establish flux, enzyme activity, or causation. Third, our bacterial communities were defined *in vitro* consortia rather than complete clinical microbiomes, only two timepoints were sampled, and the *E. faecalis* monoculture was underpowered (n = 4). Bacterial load also differed across conditions by design, as uropathogens were inoculated at lower density to offset their faster growth. The co-variation signal, however, did not track biomass, since *E. faecalis* drove disproportionate consortium-level rewiring despite a limited standalone footprint, and early differentiation was urobiome-driven even though uropathogens outgrow the community over time.

In conclusion, this study establishes that urobiome metabolic organization operates through a two-tiered architecture: a compositionally redundant temporal metabolic program that executes across single-species dropouts, and a single non-redundant keystone - *Winkia anitrata* - whose removal triggers near-total topological collapse that no other community member recapitulates. Superimposed on this architecture, pathogen invasion redirects community co-variation into a distinct emergent state: *E. coli* is the primary driver of co-variation reorganization, *E. faecalis* acts as a network facilitator whose consortium-level contribution exceeds its standalone metabolic footprint, and urobiome-pathogen copresence generates a coordinated state not recapitulated by either community in isolation. Our data suggest that in UTI, urobiome-mediated colonization resistance may rely less on the overall taxonomic composition of the community, but rather more on a limited set of non-redundant metabolic functions that can be disrupted or bypassed during pathogen invasion. This is consistent with the view that the clinical impact of a taxon may depend on its metabolic context within the community rather than on its presence alone, and underscores why taxonomic classification of individual members is a poor predictor of community-level protective function. Shifting the perspective from taxonomy to function allowed us to capture these early metabolic differences at critical stages of microbial interaction. While still preliminary, this framework may provide a useful starting point for investigating microbiomes of varying complexity through their metabolic capabilities rather than their taxonomic profiles, and for assessing how broadly such function-centered principles apply across systems.

## Materials and methods

### Experimental design

Two complementary defined microbial communities were evaluated in co-culture with iPSC urothelial barrier cultures derived from organoids. Culture conditions were maintained in line with previously described methods [Bindas et al, manuscript in preparation]. In short, iPSCs were differentiated into definitive endoderm, then urothelial lineages were expanded as organoids from a female donor line (NAS2 or EDi002-A). Dissociated organoids were seeded on 24-well transwell barriers, then for 7 days cultured in expansion medium, composed of Advanced DMEM/F12 (Fisher, 11540446) with 50% conditioned L-WRN medium (ATCC, CRL-3276), 1X B27, 1X N-2 supplement (Gibco, 11520536), 1X HEPES (Gibco, 15630056), 1X glutamax (Gibco, 35050061), 1.25 mM N-acetyl-cysteine (Sigma, A9165), 10 mM Nicotinamide (Tocris, 4106), 0.5 µM A 83-01 (Tocris, 2939), 50 ng/mL EGF (Stem Cell, 78006.1), 10 mM SB202190 (Stem Cell, 17100102), 25 ng/mL FGF7 (Novopro Labs, 318990), 100 ng/mL FGF10. 7 days of expansion medium allowed a confluent, multilayered barrier to form. After, differentiation gradient was then applied for 3 days, where expansion medium was maintained on the basolateral side of the transwell, then on the apical side of the transwell, differentiation medium was added. Differentiation medium was composed of phenol red free DMEM/F12 with L-glutamate and HEPES (Gibco, 11039021) with 5% conditioned Noggin medium kindly obtained from Dr. Vanessa Muncan and Dr. Gijs van der Brink, 5% conditioned R-spondin 1 (Trevigen, 3710-001-01), 1X B27, 1X N-2 supplement, 1.25 mM N-acetyl-cysteine, 25 ng/mL FGF7, 100 ng/mL FGF10, 1 µM PD153035, 1 µM Troglitazone, and 10 µM retinoic acid. Following this, for 3 days differentiation medium was added to the basolateral side of the transwell (850 µL) and SimUrine was added to the apical side (400 µL). Artificial, chemically-defined SimUrine was prepared per previously described methods [62].

Each bacterial strain was grown from frozen stocks either for 48-hours in a revival medium (modified NYC III with cysteine). Bacterial strains were derived from Marjon de Vos lab (University of Groningen, The Netherlands) [63], [64]. The patient-derived community strains were isolated from urine samples that were collected as part of the NWO ENW-XL project Urinary tract Infections Revisited. The lactobacilli and *G. vaginalis* were derived from the Alan Wolfe-collection (Loyola University Chicago, Maywood, IL, Unites States). *E. faecalis* and *E.coli* in the first experiments were derived from Croxall 2011 study [65], [66]. Uropathogenic *E. coli* cultures were only cultured for 24 hours due to the faster doubling time. Revival medium is composed of bacto proteose peptone no. 3 (1.5% w/v), glucose (0.5 % w/v), HEPES (0.24% w/v), NaCl (0.5 % w/v), yeast extract (0.38% w/v), inactivated newborn calf serum (5% v/v), inactivated horse serum (5% v/v). Culture in filled 15 mL tubes with cysteine was conducted to minimize oxygen levels. Bacterial suspensions were centrifuged at 3000G for 10 minutes to pellet cultures, resuspended in SimUrine, then diluted to a 0.2 OD.

Prior to inoculating transwell cultures, bacterial treatment groups were then assembled in a 10% dilution, except in the case of invading uropathogens (*E. faecalis* and uropathogenic *E. coli*) which were added at 1% to offset their markedly faster growth relative to the urobiome members, particularly UPEC. Extracellular metabolites from the apical transwell were collected at 3 h, to capture early regulatory and competitive interactions, and 6 h, to reflect downstream metabolic consequences.

After 3 hours of co-culture, as much apical SimUrine suspension as possible was removed without impacting the barrier cultures, then 400 µL of fresh SimUrine was added to each well. Apical media was then centrifuged at 3000G for 10 minutes, and the resulting supernatant was stored at -80 °C until samples were processed for untargeted metabolomics. Urothelial cultures were inspected throughout to ensure structural integrity: tight junction integrity was maintained across all experimental conditions, supporting attribution of the observed metabolic changes to microbial activity rather than to urothelial damage. No basolateral bacteria were detected in any condition.

In the infection experiment, three community configurations were studied: (i) a defined urobiome composed of *L. crispatus*, *L. gasseri*, and *G. vaginalis*; (ii) a pathogen consortium of clinically isolated uropathogenic *E. coli* and *E. faecalis* [65]; and (iii) copresence of both communities simultaneously. Single-organism controls (*E. coli* alone, *E. faecalis* alone, organoid only) were included to decompose consortium-level effects. Biological replicates per infection-model condition, at each timepoint, were host-only organoid control, n = 6; urobiome, n = 6; *E. coli* monoculture, n = 6; *E. faecalis* monoculture, n = 4; pathogen consortium, n = 4; urobiome with *E. coli*, n = 6; urobiome with *E. faecalis*, n = 4; and urobiome with the pathogen consortium, n = 4, giving 80 organoid samples in total. The design is unbalanced: two of the six independent experiments did not yield usable material for the four *E. faecalis*-containing conditions (Supplementary Table S9). In the single-strain dropout, the six-member reconstructed community isolated from a postmenopausal donor without a history of rUTI, recruited within a cohort of postmenopausal women with a history of rUTI were reassembled in vitro (*W. anitrata*; *E. faecalis* strain 1; *E. faecalis* strain 2; *S. vaginalis*; *S. pasteuri*; *S. epidermidis*). The community was studied in full and with each strain removed individually, yielding seven community conditions, plus sterile. Independent biological replicates were maintained across all conditions and timepoints (n=5).

### Metabolite extraction and LC–MS/MS acquisition

Metabolites were extracted from SimUrine supernatants using biphasic liquid–liquid extraction (2:2:1 v/v MeOH:ACN:H_2_O), protein precipitation, and stable-isotope-labelled internal standard spiking (creatinine-(N-methyl-d_3_), succinic acid-(13C_4_), L-phenylalanine-(3,3-d_2_) all purchased at Cambridge Isotope Laboratories; 5 µM final concentration; [67]). Samples were analyzed by pHILIC LC–MS/MS in positive and negative electrospray ionization modes. An enhanced data-dependent Top5 acquisition strategy [68] was applied: inclusion and exclusion lists derived from pooled quality control runs deprioritized abundant precursors with redundant fragmentation profiles, improving coverage of lower-abundance features (MS1 > 1 × 10^4^; MS2 events < 2).

### LC–MS/MS data processing, annotation, and chemical space visualization

#### Feature detection and preprocessing

Raw LC–MS data were converted to open format with MSConvert and processed using MZMine4 [25], comprising chromatographic feature detection, sample alignment, and adduct/isotope grouping. Features were filtered to remove technical noise and low-confidence signals. Complete per-run MZMine parameter sets are provided in Supplementary Methods SM1.

#### Metabolite annotation

Annotation integrated six complementary sources applied sequentially: (i) spectral library searching in MZMine 4 against public, semi-targeted, and in-house authentic standard libraries; (ii) feature-based molecular networking via GNPS2 [42]; (iii) confidence-aware molecular networking via bootstrapped edge support (SpecReBoot) [69]; (iv) in silico structure and class prediction by SIRIUS/CSI:FingerID/CANOPUS [70], [71], [72]; (v) MS2Query spectral embedding-based analog prediction [73]; and (vi) DreaMS transformer embeddings for analog search [74]. A custom consensus notebook consolidated all sources per feature into a single annotation, with MSI confidence levels assigned following Metabolomics Standards Initiative guidelines (Supplementary Figure S1) [75]. The adduct-aware feature collapsing merges redundant features sharing the same identity, summing peak areas across samples. Full annotation workflow parameters are in Supplementary Methods SM1.

#### Chemical space visualization

Joint positive- and negative-mode spectral embeddings were computed using MS2DeepScore2 [30], [76] and projected via UMAP, with features colored by CANOPUS-predicted NPC macro-class. Parameters are in Supplementary Methods SM2.

### Multivariate statistical analysis

PLS-DA was performed separately per ionization mode with biological condition as the grouping variable. Prior to PLS-DA, the feature intensity matrix was filtered to retain features present (non-zero) in at least 50% of samples, missing/zero values were imputed at one-fifth of the per-feature minimum, intensities were log2(x+1)-transformed, normalized by probabilistic quotient normalization (PQN), and Pareto-scaled. PLS-DA was fit using PLSRegression (scikit-learn, n components = 5) on a one-hot-encoded group matrix; variable importance in projection (VIP) scores were computed across all retained components. Latent variable (LV) percentages report each LV’s score variance as a proportion of total variance in the preprocessed feature matrix, as PLS components are selected to maximize covariance with class labels rather than to maximize X-variance. PERMANOVA (Bray-Curtis; 999 permutations; BH correction applied within each factor) tested the significance of community composition and timepoint. PERMDISP was applied in parallel to distinguish centroid-driven differences from within-group dispersion effects. A biological replicate was defined as an independent transwell organoid culture, inoculated and sampled separately for each condition and timepoint; replicate numbers per condition are as specified in the experimental design.

### PhenoRewire analytical framework

PhenoRewire is an interpretable, network-based framework, implemented as a Python command-line tool, designed to characterize how the co-variation structure of the metabolome is reorganized between biological states.

It is intended to complement differential abundance testing as it resolves coordinated shifts in co-regulation that abundance testing alone does not capture, including for features whose mean level is unchanged. Its outputs are intended for hypothesis generation.

#### Inputs and preprocessing

PhenoRewire accepts two tabular inputs: a feature intensity table (one row per metabolite, one column per sample) and a metadata table (sample identifiers, biological group, timepoint).

Prior to analysis, the feature intensity table was filtered by a minimum non-zero intensity threshold of 1 and by global prevalence within each run (0.50 for temporal analyses, 0.35 for phenotype analyses). Intensities were then median-normalized across samples and log2(x + 1)-transformed.

#### Feature selection

Each feature was tested for association with a binary phenotype vector (reference = 0, case = 1) or a numeric time variable using Spearman rank correlation. Statistical significance was assessed by permutation testing (1,000 permutations per feature). Empirical p-values were adjusted for multiple testing using the Benjamini–Hochberg procedure. Base FDR thresholds were 0.05 (phenotype mode) and 0.01 (temporal mode). To prevent biologically plausible comparisons from yielding empty feature sets solely due to marginal underpowering, adaptive but bounded threshold relaxation was implemented: the FDR ceiling was raised incrementally only when fewer than five features were retained, up to a maximum of 0.20, preserving interpretability while avoiding all-or-none analytical failure.

#### Network reconstruction

Following feature selection, one Spearman correlation network was reconstructed per biological state from the selected feature set, using only samples belonging to that state. Edge retention required both an absolute correlation threshold and an edge-level BH-adjusted FDR threshold: |rho| ≥ 0.70 and BH-FDR ≤ 0.05 for phenotype analyses; |rho| ≥ 0.80 and BH-FDR ≤ 0.01 for temporal analyses. Adaptive but bounded thresholding was applied when necessary to avoid networks too sparse or too dense for meaningful rewiring analysis.

#### Rewiring quantification

The edge sets of the two state-specific networks were compared systematically. Edges were classified as shared between both states, specific to state “reference”, specific to state “case”, or sign-switching (correlation sign reversal between states). Node-level rewiring scores, defined as the sum of state reference-specific and state case-specific, and sign-switching edges for each node, constituted the metabolite triage ranking, prioritizing features whose co-variation neighborhood is most restructured between states. For each state-specific network, topological descriptors were computed: degree, degree centrality, weighted degree (|rho| as weight), betweenness centrality (distance defined as 1/|rho|), eigenvector centrality, Louvain community membership, and modularity. These were computed within each real biological network rather than on the union graph, so that node importance retains direct state-specific biological meaning.

#### Outputs

Each run generates selected feature tables with group means and log2FC, state-specific network summaries, global and node-level rewiring tables, ranked metabolite triage scores, state-specific networks in GraphML format compatible with downstream spectral network integration, and final reports in Markdown and JSON format.

PhenoRewire quantifies differential co-variation structure; all outputs are associative and do not establish flux, enzyme activity, or causation. Full analysis design, per-experiment run specifications, and per-run parameter tables are provided in Supplementary Methods SM4.

### Spectral network robustness and annotation propagation via two-dimensional network visualization

PhenoRewire identifies metabolite features based on their co-variation behavior across biological states, independently of their annotation status. Because the selected feature sets frequently include low-confidence or unannotated features (MSI levels 4–5), chemical interpretation of the rewiring signal would otherwise be incomplete. To address this, features were projected onto a spectral similarity network constructed using SpecReBoot in matchms mode [69], applied to MGF files from both experiments. Bootstrap pseudo-replicates (*B* = 30) were generated by resampling discretized fragment bins; edge robustness was quantified as the proportion of replicates in which two spectra were recovered as reciprocal top-5 nearest neighbours. MS2DeepScore (k = 5, similarity threshold = 0.8, support threshold = 0.5) was selected to capture structural relationships beyond exact fragment matching, enabling edges between spectrally related features even in the absence of shared diagnostic ions. Within this spectral network, annotation was propagated from annotated features to their unannotated neighbours, allowing structural and class-level information to be transferred to features that passed Spearman correlation selection but lacked confident direct annotation. A concrete example of annotation propagation is thiocystine from the infection model experiment: the feature itself lacked a direct spectral library match, and its identity was recovered through annotation propagation in the SpecReBoot confidence-aware network, where a bootstrapped (’rescued’) edge linked it via MS2DeepScore spectral similarity to a confidently annotated neighbour [66].

The two-dimensional node-link diagrams generated from this network serve primarily as a navigational aid for annotation propagation and rapid visual triage: node size encodes the PhenoRewire rewiring score (biological relevance), and edges encode SpecReBoot-derived spectral similarity (chemical relatedness). However, the 2D layout necessarily compresses network topology, and is not intended as the primary medium for detailed inspection of rewiring structure. For in-depth exploration, including examination of edge weights, node metadata, subnetwork topology, and community structure, the full GraphML output should be loaded directly into Cytoscape, which preserves the complete graph structure and supports interactive filtering, custom visual mappings, and integration with annotation databases. The 2D diagrams were generated using a custom Python layout pipeline; a base implementation, designed to accept any GraphML input (of which SpecReBoot-derived networks are an example), is available in the PhenoRewire repository.

### Comparative genomics of community isolates

Genome assemblies were available and kindly provided by dr. Marjon de Vos for the six bacterial isolates comprising the synthetic urobiome (*W. anitrata*, *S. vaginalis*, two *Enterococcus faecalis* isolates - named *E. faecalis* strain 1 and *E. faecalis* strain 2, *S. pasteuri*, and *S. epidermidis*). Genome-based taxonomic resolution of the two *E. faecalis* isolates was performed using the Type (Strain) Genome Server (TYGS), with pairwise digital DNA-DNA hybridization (dDDH) estimated under the d4 formula, which is appropriate for draft-quality genomes.

Predicted protein sequences were functionally annotated with eggNOG-mapper, assigning each protein to eggNOG orthologous groups and Clusters of Orthologous Genes (COG) categories; outputs were parsed in Python/pandas and COG identifiers extracted by regular-expression matching of the COG numerical-identifier pattern. For each isolate, the total number of annotated proteins, the number of proteins with a COG assignment, COG coverage (%), and the number of distinct COG identifiers were computed; COG coverage ranged from 83.70% to 89.32% across the six isolates. A binary COG presence/absence matrix was constructed across all six isolates, and COGs were classified as conserved (present in all six), shared (present in two to five), or isolate-specific (present in one). Pairwise COG-content similarity was assessed using the Jaccard index (SciPy) on the binary matrix and visualized as a lower-triangular heatmap, and global isolate relationships were summarized by principal coordinates analysis (PCoA) on the corresponding Jaccard distance matrix. The two *E. faecalis* isolates were additionally compared directly to identify COGs present in one isolate but absent in the other.

### Host-microbiome metabolic crosstalk annotation

To characterize host-microbiome metabolic crosstalk, PhenoRewire was applied to 21 comparisons spanning both ionization modes and two experimental axes: phenotype (community composition) and time (3 h vs 6 h). Co-variation networks were built only from features that were differentially abundant between conditions.

Each differentially abundant feature was further classified by whether its network connectivity also changed between conditions. Differentially abundant and rewired features changed in both abundance and connectivity; differentially abundant features changed in abundance only. Connectivity change (rewiring) was quantified by counting single-state edges: network connections present in only one condition and absent from the comparator. This captures changes in feature co-variation that a standard abundance test cannot detect. Reproducibility across comparisons was assessed by ranking metabolites by (i) the number of comparisons in which they were rewired, and (ii) whether the effect was concordant between positive and negative ionization mode.

Four metabolic axes were tracked as targeted readouts across all comparisons: the arginine deiminase (ADI) pathway (arginine, citrulline, ornithine, N-carbamoyl derivatives, and ureidosuccinic acid), the allantoin/purine-catabolism axis (allantoin, uric acid, and urea), tryptophan, and proline.

For the strain-dropout experiment, each rewired feature was attributed to a single community member by leave-one-out (LOO) analysis: the member whose removal caused the largest drop in that feature’s rewiring score, relative to the full community, was assigned as the attributed strain. Features significant against the sterile control in the full community, but not in any single-strain dropout, were classified as community-emergent.

Because the experiment did not include the sterile arm, host and microbial contributions to crosstalk could not be decomposed additively; all crosstalk inference therefore derives from differential network connectivity between co-culture conditions.

Networks with modularity Q < 0.3 were flagged as low-confidence, and module-level interpretation was withheld for these partitions.

## Data availability

Raw metabolomics data will be deposited in the MetaboLights repository (EMBL-EBI) and released upon peer-reviewed publication of this work. Accession numbers will be added to this statement in a revised version of this preprint. The endogenous metabolite spectral library used for this study is available at Zenodo (https://zenodo.org/records/21394919). Genome assemblies for the six bacterial isolates used in the strain-dropout experiment are available at ENA Project PRJEB124534.

## Code availability

PhenoRewire version 0.1.0 is freely available at https://github.com/ldellavedova/PhenoRewire. All other software used in this study is publicly available and cited with version numbers in the Methods (MZmine 4, SIRIUS/CSI:FingerID/CANOPUS, GNPS2, MS2Query, DreaMS, MS2DeepScore, SpecReBoot).

## Supporting information

Supplementary Information contains a glossary of terms, Supplementary Figures S1-S6, Supplementary Tables S1-S9, and Supplementary Methods SM1-SM4. Supplementary Tables S1-S6 and S8 are additionally provided as separate Excel files.

## Author contributions

L.D.V., C.R.B. and J.J.J.v.d.H. conceived and designed the metabolomics study and analytical framework. A.J.B. and J.M.W. conceived and designed the urothelial organoid and bacterial co-culture experiments. L.D.V. developed the PhenoRewire framework and computational pipeline, performed the metabolomics data analysis, and wrote the original draft. A.J.B. performed the urothelial organoid and co-culture experiments and contributed to writing. J.J. contributed to mass spectrometry method setup and data acquisition. M.T.D. performed metabolomics experiments and contributed to metabolomics data analysis. M.T.D. and D.G.-V. performed comparative genomics analysis of community isolates. P.G.M. prepared the SimUrine medium and preparation for sequencing. J.K.B. performed isolate sequencing. T.H. performed genome assembly and bioinformatics of the sequenced isolates. Z.F. contributed to cell expansion and bacterial culturing. A.F. contributed to sample collection. J.M.W. supervised the host–microbe interaction work. M.G.J.d.V. provided the bacterial isolates and synthetic community and supervised the microbial ecology work. C.R.B. supervised the metabolomics work and the project. J.J.J.v.d.H. supervised the computational metabolomics work and the project. C.R.B., M.d.V. and J.J.J.v.d.H. reviewed and edited the manuscript. J.M.W., M.G.J.d.V., C.R.B. and J.J.J.v.d.H. acquired funding. All authors reviewed and approved the manuscript.

## Supporting information

Supplementary_Table_S1_infection_model

Supplementary_Table_S2_infection_model_PLSDA_VIP

Supplementary_Table_S3_infection_model_network_stats

Supplementary_Table_S4_strain_dropout_model

Supplementary_Table_S5_strain_dropout_PLSDA_VIP

Supplementary_Table_S6_strain_dropout_network_stats

Supplementary_Table_S8_DA_vs_rewiring_benchmark

## Acknowledgments

We thank Alan Wolfe for the use of some isolates derived from his collection and for the revival medium protocol. We thank Oshin Vellalara for the support in endpoint measurements and sample collection. We would like to thank the Urinary tract infections revisited (UTIr) consortium for the fruitful discussion and feedback.

