## Supplementary_Information for "Metabolite co-variation networks reveal keystone functions and an emergent pathogen state in the human urobiome"

### Glossary

| Term | Meaning in this study |
|---|---|
| rUTI | Recurrent urinary tract infection. |
| iPSC | induced pluripotent stem cells. |
| dDDH | Digital DNA–DNA hybridization: a genome-sequence-based estimate of how closely two isolates are related. Values above roughly 70% indicate the same species. |
| COG / eggNOG | Clusters of Orthologous Genes: functional categories assigned to predicted proteins, here via the eggNOG database. |
| Jaccard index | Similarity between two presence/absence lists: shared items divided by total distinct items. 1 = identical, 0 = no overlap. |
| PCoA | Principal coordinates analysis |

| | |
|---|---|
| Untargeted metabolomics | Measurement of all detectable small molecules in a sample without a predefined list of compounds to look for. |
| Feature | One detected ion, defined by its mass-to-charge ratio and retention time. A feature is not necessarily an identified compound, and one compound can give rise to several features. |
| *m/z* | Mass-to-charge ratio: what the mass spectrometer actually measures. |
| Ionization mode (positive / negative) | Molecules must carry a charge to be detected. Positive and negative mode favour different chemistries, so the two modes give complementary, partly overlapping coverage of the metabolome and are analysed separately throughout. |
| pHILIC | A chromatography column chemistry that retains and separates polar metabolites, which conventional reversed-phase columns do not. |
| MS1 / MS2 | MS1 is the survey scan that records intact ions; MS2 is the fragmentation spectrum of a selected ion, which carries the structural information used for identification. |
| Data-dependent acquisition (Top5) | The instrument automatically fragments the five most intense ions in each cycle. |
| Adduct | The charged form in which a molecule is detected, for example $[M+H]^+$ or $[M-H]^-$. Adduct-aware collapsing merges features that are different adducts of the same compound. |
| Spectral library matching | Comparing a measured MS2 spectrum against reference spectra of known compounds. |
| Molecular networking (MN) | Connecting features whose MS2 spectra are similar, so that structural analogues form clusters even when only some members are identified. |
| Annotation propagation | Transferring a compound name or class from an annotated feature to an unannotated feature that is its close spectral neighbour. |
| MSI confidence level | A 1–5 scale for how certain an annotation is: 1, matched to an authentic standard; 2, probable library match; 3, putative — either a compound class predicted *in silico*, or a tentative compound name whose measured *m/z* could not be reconciled with a common adduct of the corresponding formula; 4, molecular formula only; 5, unknown. |

| | |
|---|---|
| UMAP | A dimensionality-reduction method used to lay out many spectra on a two-dimensional map so that chemically similar ones fall close together. Positions are qualitative: proximity is meaningful, absolute axis values are not. |

| | |
|---|---|
| PLS-DA | Partial least squares discriminant analysis: a supervised model that searches for the combinations of metabolites that best separate groups the experimenter has defined in advance. |
| Latent variable (LV) | A composite axis built by PLS-DA from many metabolites at once. The percentage reported for each LV is the share of total variance in the preprocessed feature matrix that the axis captures. |
| Score plot | The samples drawn in the space of the latent variables. Samples that sit close together have similar metabolic profiles. |
| Loading | The weight of one metabolite on one latent variable: how much that metabolite contributes to that axis, and in which direction. |
| VIP score | Variable Importance in Projection: a single number summarising how much a metabolite contributes to the group separation across all components of the PLS-DA model. |
| Q2 | A cross-validated measure of how well the model predicts samples it was not trained on. It guards against a model that separates groups only because it has memorised the data. |
| PQN | Probabilistic quotient normalisation: corrects for samples being globally more or less concentrated. |
| Pareto scaling | Scaling each metabolite by the square root of its standard deviation. |
| PERMANOVA | A permutation-based test of whether the multivariate centroids of two or more groups of samples differ. The pseudo-F statistic expresses the size of the effect: larger means the groups are further apart relative to their internal scatter. |
| PERMDISP | The companion test to PERMANOVA, asking whether groups differ in internal spread. |
| Bray–Curtis dissimilarity | An abundance-weighted measure of how different two samples are: 0 means identical, 1 means no shared signal. |
| Permutation test / empirical P | Significance obtained by repeatedly reshuffling the sample labels and asking how often chance reproduces the observed result. |
| FDR / q value | False discovery rate. The Benjamini–Hochberg q value is the expected proportion of false positives among the results declared significant. |
| Spearman rho | A rank-based correlation coefficient between −1 and +1. |

| **Term** | **Meaning in this study** |
|---|---|
| Co-variation | Two metabolites rising and falling together across samples. |
| Co-variation network | A graph in which nodes are metabolites and edges join metabolite pairs whose correlation passes both a strength and a significance threshold. |
| Node / edge | A node is one metabolite; an edge is one retained correlation between two metabolites. |
| Rewiring | A change in which metabolites are correlated with which, between two states. This is the central readout of PhenoRewire: a metabolite can be |

| | |
|---|---|
| | strongly rewired while its average level does not change at all. |
| Shared / state-specific | An edge present in both states, or present in only one, or present in both but with the correlation reversing sign. |
| Rewiring score | Per metabolite, the number of state-specific plus sign-switching edges it carries. Used to rank metabolites. |
| Network density / edge count | Measures how interconnected a network is. A denser late network means more metabolites moving in a coordinated way. |
| Modularity (Q) | How cleanly a network divides into separate modules. Values near 0 indicate near-random wiring with no distinguishable groups; higher values indicate well-separated modules. Below Q = 0.3 the module boundaries are not trustworthy. |
| Louvain community detection | The algorithm used to partition a network into modules and from which modularity is computed. |
| Degree / betweenness / eigenvector centrality | Three complementary measures of a metabolite's position: how many partners it has, how often it lies on paths between other metabolites, and how strongly it connects to other well-connected metabolites. |
| Phenotype mode / temporal mode | The two ways PhenoRewire can be run: comparing two community compositions at one timepoint, or comparing one composition between two timepoints. |
| LOO attribution | Assigning a rewired metabolite to the community member whose removal most reduces that metabolite's rewiring score relative to the full community. |
| GraphML | The file format in which the networks are exported, readable by Cytoscape for interactive inspection. |

**Supplementary Figures**

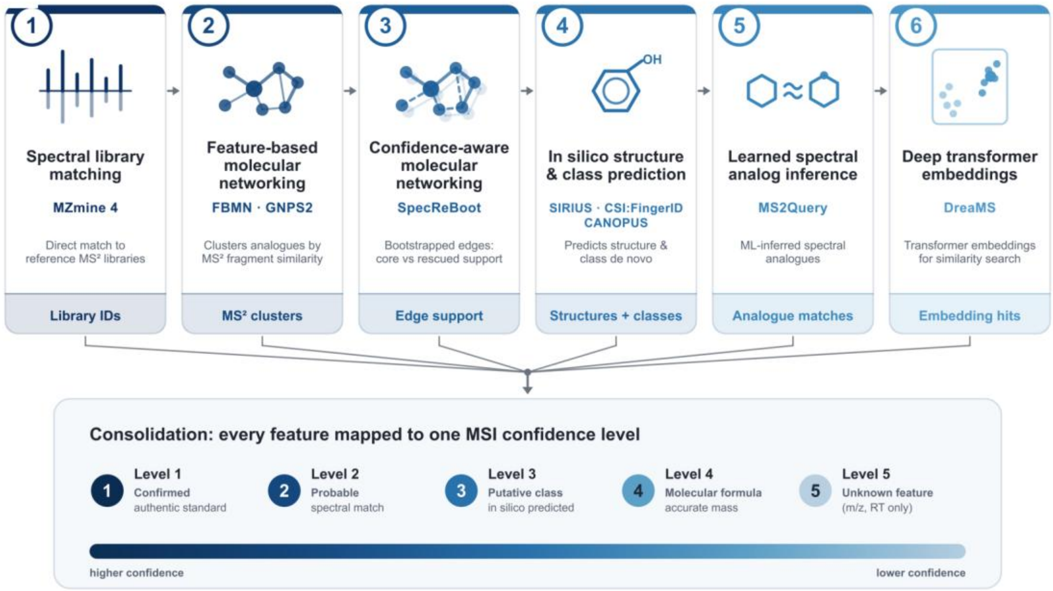


**Supplementary Figure S1. The six-tool integrative metabolite annotation pipeline.** Untargeted LC–MS/MS features were annotated by six complementary approaches applied in sequence: (1) spectral library matching against public, semi-targeted and in-house authentic-standard libraries (MZmine 4); (2) feature-based molecular networking (FBMN/GNPS2); (3) confidence-aware molecular networking via bootstrapped edge support (SpecReBoot); (4) *in silico* structure and compound-class prediction (SIRIUS/CSI:FingerID/CANOPUS); (5) learned spectral analogue inference (MS2Query); and (6) deep transformer-based embedding search of analogues (DreaMS). Per-feature evidence from all six sources was consolidated by a custom consensus notebook into a single annotation, and every feature was assigned one Metabolomics Standards Initiative (MSI) confidence level: 1, confirmed against an authentic standard; 2, probable spectral/library match; 3, putative (i.e. either a compound class predicted *in silico*, or a tentative compound name retained for information but not confirmed); 4, molecular formula from accurate mass; 5, unknown feature (*m/z* and retention time only). Adduct-aware feature collapsing then merged redundant features sharing the same identity, summing peak areas across samples. Full per-tool parameters are given in Supplementary Methods SM1.

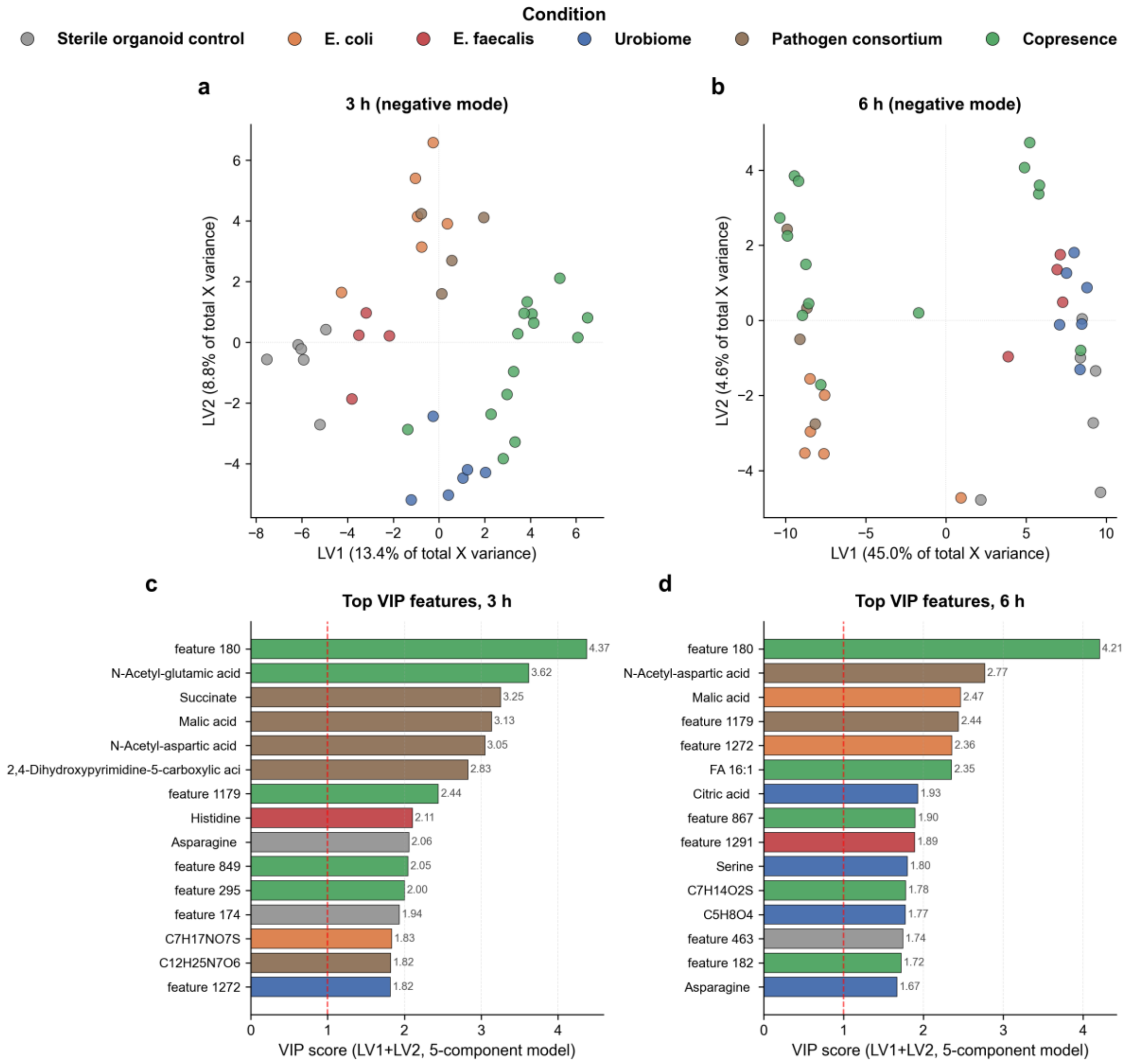


**Supplementary Figure S2. PLS-DA of the infection model in negative ionization mode reproduces the temporal shift in discriminant metabolite identity seen in positive mode.** (**a**,**b**) PLS-DA score plots (negative ion mode) at 3 h (**a**) and 6 h (**b**), with samples coloured by experimental condition. Axis labels report each latent variable's (LV) score variance as a percentage of total variance in the preprocessed feature matrix. At 3 h, urobiome-containing samples separate from all other conditions along LV2, while *E. coli*- and pathogen-consortium-containing samples occupy a separate area os the score plot. By 6 h the axis roles converge with the positive-mode result: *E. coli*-containing conditions (*E. coli* monoculture, pathogen consortium, copresence) separate along LV1, while urobiome and host-only (sterile) samples converge. (**c**,**d**) Top 15 discriminant features ranked by Variable Importance in Projection (VIP) score at 3 h (**c**) and 6 h (**d**); bar colour indicates the condition with the highest mean intensity for that feature and the dashed line marks the VIP = 1.0 threshold. At 3 h the highest-ranked features are dominated by TCA-cycle and amino-acid-derivative species (succinate, malic acid, N-acetyl-glutamic acid, N-acetyl-aspartic acid); by 6 h the signature broadens to include additional TCA intermediates (citric acid), fatty acids and proteinogenic amino acids (serine, asparagine), mirroring the temporal broadening described for positive mode in main text.

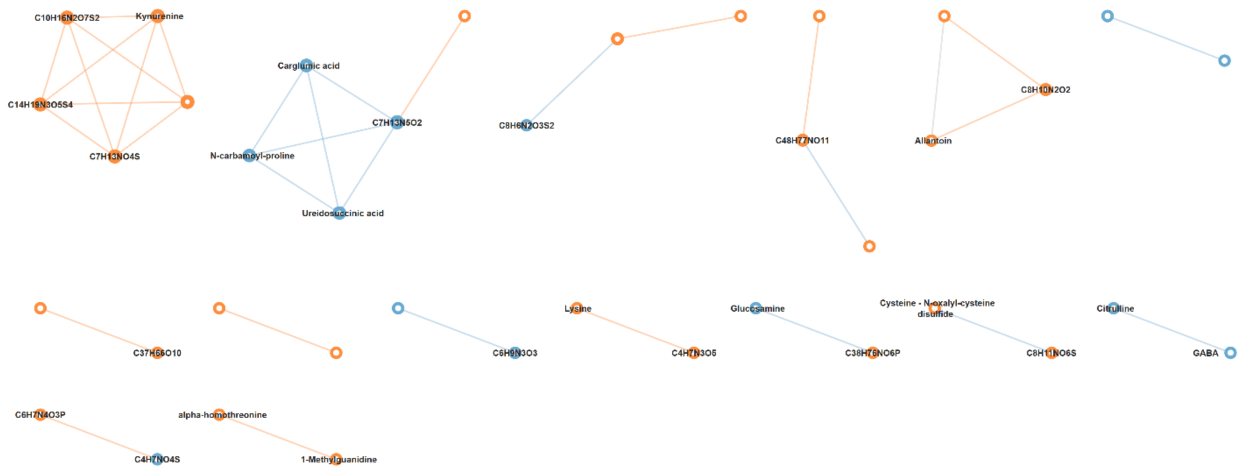


**Supplementary Figure S3. PhenoRewire temporal co-variation network of the urobiome condition, positive ionization mode.** Temporal rewired network of the metabolite features significantly associated with the 3 h to 6 h transition in the defined urobiome. Node colour encodes timepoint association (blue, 3 h; orange, 6 h) and edges are coloured by state specificity (blue, 3 h only; orange, 6 h only; grey, shared). The network remains sparse at both timepoints. The 3 h state is characterised by the carbamoyl-phosphate/nitrogen signature (citrulline, ureidosuccinic acid, N-carbamoyl-proline, carglumic acid / N-carbamyl-glutamic acid), and the 6 h state extends only modestly towards kynurenine, cysteine-N-oxalyl-cysteine disulfide and alpha-homothreonine, without the large-scale edge expansion seen in the pathogen and copresence conditions showed in the main text. Edge counts and modularity for all conditions are given in Supplementary Table S3.

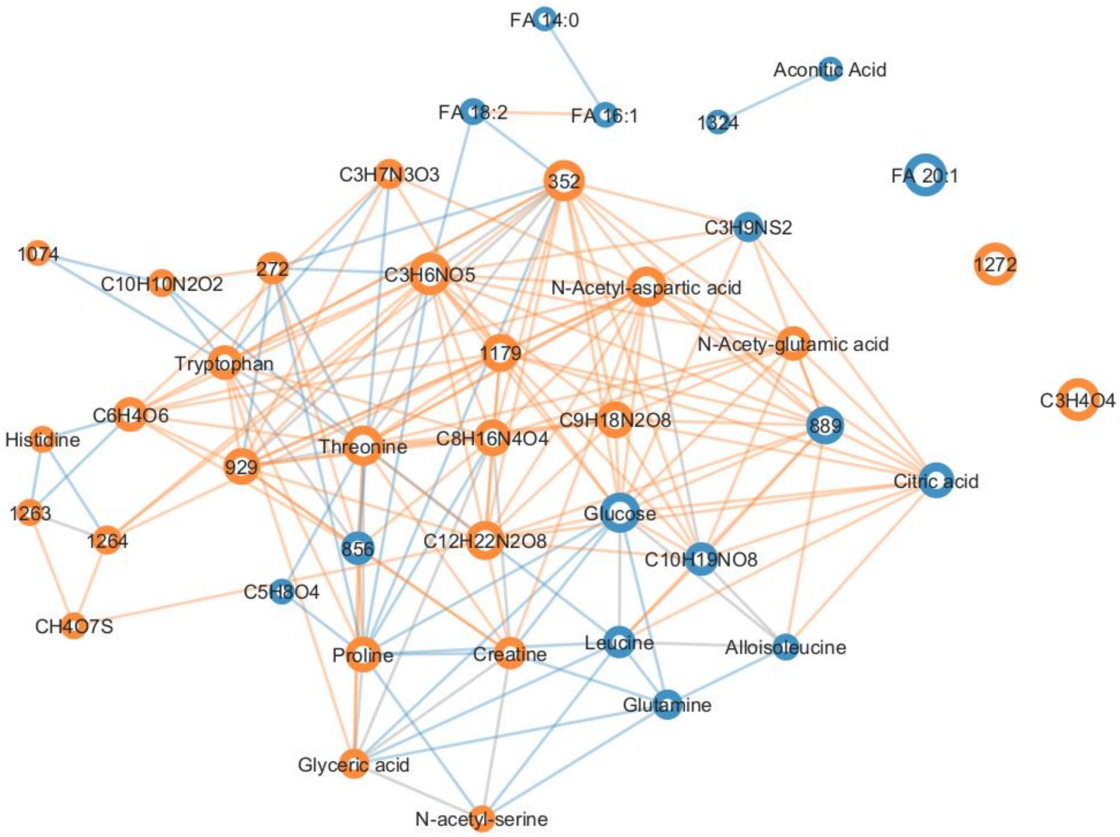


**Supplementary Figure S4. PhenoRewire temporal co-variation network of the pathogen phenotype, negative ionization mode.** Temporal co-variation network for the pathogen consortium between 3 h and 6 h in negative ionization mode. The directional shift from early citric acid, glucose, glutamine co-variation to late histidine, threonine, proline, tryptophan, creatine co-variation is reproduced across polarities, confirming cross-mode consistency of the temporal metabolic transition described in the main text for positive mode. Node and edge encoding as in Supplementary Figure S3.

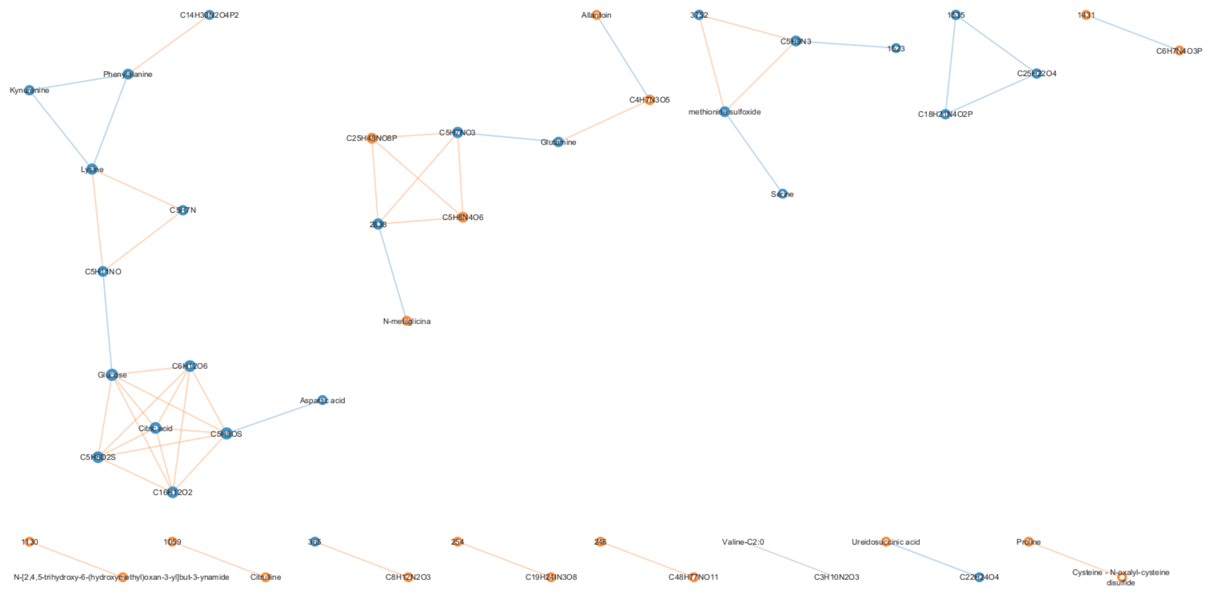


**Supplementary Figure S5. PhenoRewire temporal co-variation network of the *E. coli* monoculture, positive ionization mode.** Temporal co-variation network for the *E. coli* monoculture between 3 h and 6 h. The *E. coli* monoculture reproduces the directional co-variation architecture of the pathogen consortium at reduced topological magnitude, identifying *E. coli* as the primary architect of the pathogen temporal signal. The *E. faecalis* monoculture is not shown because it yielded no significant temporal features in either polarity and no differentially abundant features at either timepoint; its standalone co-variation architecture was indistinguishable from the host-only background. Node and edge encoding as in Supplementary Figure S3.

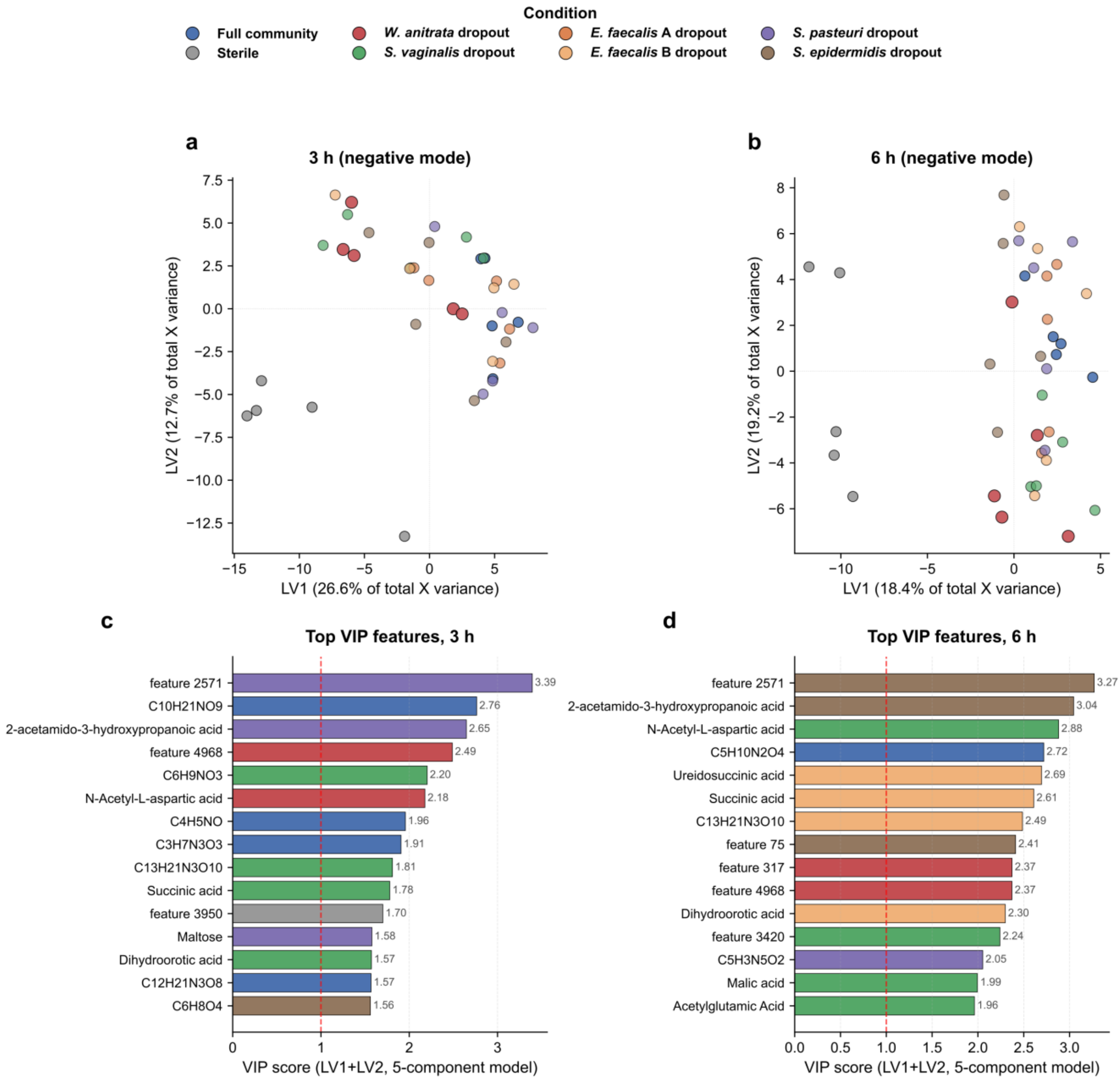


**Supplementary Figure S6. PLS-DA scores and VIP discriminants of the strain-dropout model, negative ionization mode.** (**a**,**b**) Score plots at 3 h (**a**; LV1 = 26.6%, LV2 = 12.7% of total X variance) and 6 h (**b**; LV1 = 18.4%, LV2 = 19.2%); each latent variable is expressed as a percentage of total X variance. Samples are coloured by community configuration (n = 5 per condition per timepoint). The sterile control separates from all bacterial conditions at both timepoints. Among the bacterial configurations, the *W. anitrata* dropout is the most displaced from the full-community centroid at 3 h, although within-group spread is of comparable magnitude, so this is a shift rather than a discrete cluster. By 6 h all dropouts contract while residual compositional structure persists, consistent with the negative-mode PERMANOVA signal retained at 6 h. (**c**,**d**) The fifteen highest-VIP features at 3 h (**c**) and 6 h (**d**), computed from all five latent variables; bars are coloured by the condition in which the feature reaches its highest mean intensity and the dashed line marks the conventional VIP = 1 threshold. The top discriminant at both timepoints is feature 2571 (VIP = 3.39 at 3 h; VIP = 3.27 at 6 h), which is unannotated: it is reported at MSI level 5 and labelled by its row ID, as are all other features here that lack a consensus annotation.

**Supplementary Tables**

**Supplementary Table S1. PERMANOVA and PERMDISP results, infection model.**

Multivariate tests of community composition (PERMANOVA) and within-group dispersion (PERMDISP) for the infection model. Every comparison was run on three distance metrics, Bray-Curtis, Canberra and Euclidean, and the table is organised in four sections. Sections A and B report the positive ionization mode and sections C and D the negative mode. Sections A and C list pairwise comparisons, both condition pairs within a timepoint and 3 h versus 6 h within a condition. Sections B and D report the stratified global group effect across all eight conditions within each timepoint, using 999 permutations. Every section gives sample and feature counts, PERMANOVA pseudo-F with raw P and Benjamini-Hochberg q, and the parallel PERMDISP statistic, so that a significant PERMANOVA result can be checked against a dispersion artefact. Provided as a separate Excel file: Tables/Supplementary_Table_S1_infection_model.xlsx

**Supplementary Table S2. PLS-DA VIP scores, infection model.**

Section A gives PLS-DA fit statistics by ionization mode and timepoint (sample and feature counts, number of components, LV1/LV2 variance explained as a percentage of total X variance, and leave-one-out cross-validated Q2). Section B lists every feature with VIP > 1.0 at 3 h and 6 h in both ionization modes, ranked by VIP within each mode × timepoint block, with loadings and per-condition mean intensities. Provided as a separate Excel file: **Tables/Supplementary_Table_S2_infection_model_PLSDA_VIP.xlsx**

**Supplementary Table S3. PhenoRewire network statistics, infection model.**

State-specific node and edge counts, Louvain modularity and community counts, and the number of selected features for every partition of the infection model (urobiome, pathogen consortium, *E. coli* monoculture, *E. faecalis* monoculture, copresence and host-only) at 3 h and 6 h, in positive and negative ionization mode. Provided as a separate Excel file: **Tables/Supplementary_Table_S3_infection_model_network_stats.xlsx**

**Supplementary Table S4. PERMANOVA and PERMDISP results, strain-dropout model.**

Multivariate tests of community composition (PERMANOVA) and within-group dispersion (PERMDISP) for the strain-dropout model, in the same four-section layout as Supplementary Table S1 and likewise run on Bray-Curtis, Canberra and Euclidean distances. Sections A and B report the positive ionization mode and sections C and D the negative mode; sections A and C list pairwise comparisons, both condition pairs within a timepoint and 3 h versus 6 h within a condition, and sections B and D report the stratified global group effect across all eight compositional conditions within each timepoint, using 999 permutations. Column definitions are as in Supplementary Table S1. Provided as a separate Excel file: Tables/Supplementary_Table_S4_strain_dropout_model.xlsx

**Supplementary Table S5 | PLS-DA VIP scores, strain-dropout model.**

Section A gives PLS-DA fit statistics by ionization mode and timepoint; Section B lists every feature with VIP > 1.0 at 3 h and 6 h in both modes, ranked by VIP within each mode × timepoint block, with LV1/LV2 variance explained, loading values and leave-one-out Q2. Provided as a separate Excel file: **Tables/Supplementary_Table_S5_strain_dropout_PLSDA_VIP.xlsx**

**Supplementary Table S6 | PhenoRewire network statistics, strain-dropout model.**

Edge counts, Louvain modularity and number of selected features for all seven compositional conditions at both timepoints, and for all twelve phenotype-mode comparisons (full community versus each single-strain dropout, positive mode). Confirms the *W. anitrata*-absent community (Q = 0.038 at 3 h) as a categorical outlier relative to the full community (Q = 0.707) and to all five other dropout communities, and records the zero-feature outcome in eleven of the twelve phenotype comparisons. Provided as a separate Excel file: **Tables/Supplementary_Table_S6_strain_dropout_network_stats.xlsx**

**Supplementary Table S7. Bacterial strains used in this study.**

| Strain (genome-based) | Symbol | Experiment | Isolation source | Reference |
|---|---|---|---|---|
| *Escherichia coli* (uropathogenic) | E | Infection model | UTI patient (clinical isolate) | Croxall et al. 2011 [64], [65] |
| *Enterococcus faecalis* (pathogenic strain) | F | Infection model | UTI patient (clinical isolate) | Croxall et al. 2011 [64], [65] |
| *Lactobacillus crispatus* | A | Infection model | A. J. Wolfe collection | - |
| *Lactobacillus gasseri* | B | Infection model | A. J. Wolfe collection | - |
| *Gardnerella vaginalis* | C | Infection model | A. J. Wolfe collection | - |
| *Winkia anitrata* (38-M0-1) | 1 | Strain-dropout model | Postmenopausal donor without personal rUTI history (UTIr cohort) | [62], [63] |
| *Streptococcus vaginalis* (38-M0-2) | 2 | Strain-dropout model | Postmenopausal donor without personal rUTI history (UTIr cohort) | [62], [63] |
| *Enterococcus faecalis* strain 1 (38-M0-3, isolate A) | 3 | Strain-dropout model | Postmenopausal donor without personal rUTI history (UTIr cohort) | [62], [63] |
| *Enterococcus faecalis* strain 2 (38-M0-4, isolate B) | 4 | Strain-dropout model | Postmenopausal donor without personal rUTI history (UTIr cohort) | [62], [63] |
| *Staphylococcus pasteuri* (38-M0-5) | 5 | Strain-dropout model | Postmenopausal donor without personal rUTI history (UTIr cohort) | [62], [63] |
| *Staphylococcus epidermidis* (38-M0-6) | 6 | Strain-dropout model | Postmenopausal donor without personal rUTI history (UTIr cohort) | [62], [63] |

Taxonomic note: two strain-dropout isolates were reclassified following genome-based analysis. Earlier pipeline outputs and figure labels may carry the previous names *Winkia neuii* for *Winkia anitrata* and *Streptococcus anginosus* for *Streptococcus vaginalis*.

**Supplementary Table S8. Abundance-association testing versus co-variation rewiring.**

Per experiment, ionization mode and analysis axis: number of comparisons run, comparisons yielding a co-variation network, features significant by abundance association ($q < 0.05$), rewired features, and shared versus state-specific edge counts. Quantifies what co-variation analysis adds over abundance testing. Provided as a separate Excel file: **Tables/Supplementary_Table_S8_DA_vs_rewiring_benchmark.xlsx**

**Supplementary Table S9. Sample sizes and feature counts.**

A biological replicate is an independent transwell organoid culture, inoculated and sampled separately for each condition and timepoint. Replicate numbers were counted from the sample columns present in the processed feature tables and are identical in positive and negative ionization mode. The infection model comprises six independent experiments and is an unbalanced design: two of the six did not yield usable material for the four *E. faecalis*-containing conditions, which therefore carry n = 4 rather than n = 6. All analyses were run on the replicates actually available and no values were imputed to balance the design. Three further columns (SIM_01 to SIM_03) are present in the infection-model feature tables; they are not organoid biological replicates and are not included in the counts below. Strain-dropout model: n = 5 per condition across eight compositional conditions at both timepoints (80 samples), a balanced design. All samples were analysed in both positive and negative HESI modes. Raw features are the aligned MZmine 4 feature counts; preprocessed features are those retained after prevalence filtering, normalisation and transformation as described in the main text Methods.

| **Infection model condition** | **n at 3 h** | **n at 6 h** | **Samples** |
|---|---|---|---|
| Host-only (sterile) organoid control | 6 | 6 | 12 |
| Urobiome (*L. gasseri*, *L. crispatus*, *G. vaginalis*) | 6 | 6 | 12 |
| *E. coli* monoculture | 6 | 6 | 12 |
| *E. faecalis* monoculture | 4 | 4 | 8 |
| Pathogen consortium (*E. coli* + *E. faecalis*) | 4 | 4 | 8 |
| Urobiome + *E. coli* | 6 | 6 | 12 |
| Urobiome + *E. faecalis* | 4 | 4 | 8 |
| Urobiome + pathogen consortium | 4 | 4 | 8 |
| **Total organoid samples** | **40** | **40** | **80** |

| **Experiment** | **Ionization mode** | **Raw features** | **Preprocessed features** |
|---|---|---|---|
| Infection model | Positive | 1,125 | 699 |
| Infection model | Negative | 449 | 323 |
| Strain-dropout model | Positive | 4,151 | 1,410 |
| Strain-dropout model | Negative | 2,161 | 778 |

**Supplementary Methods**

**Supplementary Methods SM1. Annotation workflow parameters.**

Parameters for the six-tool integrative annotation pipeline of Supplementary Figure S1. Feature detection and preprocessing were carried out in **MZmine 4.7.28** from an archived batch file; the settings below were identical for positive and negative ionization mode and for both experiments. All m/z tolerances are given in ppm.

**SM1a. MZmine 4.7.28 feature detection and alignment**

| **Term** | **Meaning in this study** |
|---|---|
| Mass detection, MS1 | Centroided spectra; noise level $1.0 \times 10^4$, noise factor 8; fragment scans denormalised (trap). |
| Mass detection, MS2 | Centroided spectra; noise level $1.0 \times 10^4$, noise factor 4. |
| Chromatogram building (ADAP) | Minimum 8 consecutive scans; minimum intensity for consecutive scans $5.0 \times 10^4$; minimum absolute height $9.0 \times 10^4$; scan-to-scan m/z tolerance 5 ppm. |
| Smoothing | Savitzky–Golay over retention time, window 11 scans. |
| Feature resolving (local minimum search) | Chromatographic threshold 0.9; minimum search range 0.2 min; minimum absolute height $9.0 \times 10^4$; peak-top/edge ratio 1.4; minimum 8 data points. MS2 pairing: RT filter 0.25 (feature edges), minimum relative feature height 0.25, MS1-MS2 precursor tolerance 10 ppm. |
| Isotope grouping | RT tolerance 3.0; m/z tolerance 3 ppm; monotonic shape required; maximum charge 2; most intense isotope kept as representative; features carrying MS2 never removed. |
| Isotope finder | Elements H, C, N, O, S; maximum isotope charge 1; searched in the single most intense scan (feature-to-scan tolerance 3 ppm). |
| Alignment (join aligner) | m/z tolerance 5 ppm (weight 3.0); RT tolerance 4.0 (weight 1.0); same charge state not required; isotope-pattern and spectral-similarity comparison disabled. |
| Row filtering | $^{13}C$ isotope pattern validated and $^{13}C$ rows removed (maximum charge 2, minimum carbon estimated, O excluded); m/z tolerance 3 ppm; rows with MS2 and annotated rows never removed. |
| Gap filling | Multi-thread peak finder; intensity tolerance 0.2; RT tolerance 3.0; m/z tolerance 10 ppm; minimum 8 data points. |
| Duplicate filtering | New-average mode; RT tolerance 0.17; m/z tolerance 1.5 ppm. |
| Correlation grouping | RT tolerance 4.0; intensity threshold for correlation 5.0; minimum 60% intensity overlap. Feature-shape correlation: Pearson, minimum r = 0.85, minimum 5 data points (2 on the edge). Feature-height correlation: Pearson, minimum r = 0.70, minimum 2 samples. |
| Ion identity networking | m/z tolerance 3 ppm; maximum charge 2; maximum 2 molecules per cluster; annotation refinement on, networks without a monomer deleted, link threshold 4. |

Other Software versions used: MS2Query 1.5.3, MS2DeepScore 2.6.0 for the spectral embeddings and 2.0.0 within MS2Query, matchms 0.31.0 and 0.26.4 respectively, spec2vec 0.9.1 and 0.8.0 respectively, Python 3.12 and 3.9 respectively.

**Supplementary Methods SM2. Spectral embedding and UMAP projection parameters.**

Parameters for the MS2DeepScore2 embeddings and UMAP projections shown in main text Fig. 1d and Fig. 3b. Positive- and negative-mode MGF files exported from MZmine were embedded jointly with a single cross-polarity MS2DeepScore2 model (ms2deepscoremodel.pt; MS2DeepScore 2.6), so that features from the two polarities occupy one shared space. Spectra with fewer than 5 fragment peaks were excluded. Pairwise similarity was computed as the cosine similarity between embedding vectors. For the accompanying spectral network, edges were retained at a minimum similarity of 0.85, restricted to mutual nearest neighbours and capped at 10 mutual neighbours per node. The two-dimensional projection used UMAP with nneighbors = 50, mindist = 0.10, metric = cosine and randomstate = 42. Colour assignments follow CANOPUS-predicted NPC pathway macro-classes; positions are qualitative, so proximity is meaningful but absolute axis values are not.

**Supplementary Methods SM3. SpecReBoot parameters.**

SpecReBoot was run in matchms mode (specreboot matchms; Python 3.12, matchms 0.31.0, MS2DeepScore 2.6) on the MGF exports from Mzmine of both experiments, separately per ionization mode. MS2 spectra were discretized onto a global m/z bin grid rounded to two decimals (0.01 m/z bins), and bootstrap pseudo-replicates (B = 30) were generated by resampling those bins. Edge robustness was quantified as the proportion of replicates in which two spectra were recovered as reciprocal nearest neighbours among the k = 5 most similar spectra, and edges were retained at a support threshold of 0.5. The peak-matching tolerance was 0.02 Da. Similarity thresholds were 0.7 for cosine and modified cosine and 0.8 for MS2DeepScore. Remaining settings were left at their defaults. Within each network, annotations were propagated from annotated features to their unannotated neighbours, allowing structural and class-level information to reach features selected by PhenoRewire that lacked a confident direct annotation (*i.e.*, thiocystine in the infection model). Mass-shift matching during propagation used tolerances of ±15 mDa for adduct switches, ±5 mDa for isotopologues and ±5 mDa for structural modifications.

**Supplementary Methods SM4. PhenoRewire run specifications.**

Per-experiment YAML configuration files, partition definitions and per-run parameter tables for all PhenoRewire analyses reported in this study. Feature prevalence thresholds 0.50 (temporal) and 0.35 (phenotype), minimum non-zero intensity 1, and 1,000 permutations per feature. Feature selection used a Benjamini–Hochberg FDR of 0.05, relaxed automatically to a ceiling of 0.10 when fewer than five features passed. Networks were built from Spearman correlations with base thresholds of $|\rho| \geq 0.70$ (phenotype) or $\geq 0.80$ (temporal) and base edge-FDR of 0.05 (phenotype) or 0.01 (temporal); when a partition yielded fewer than three edges, the correlation threshold was relaxed towards a floor of $|\rho| = 0.50$ and the edge-FDR towards a ceiling of 0.20. Relaxation is bounded: the run is reported as failed, not relaxed further, if fewer than five features, ten edges or five nodes survive. Every relaxation event is recorded per partition in the deposited logs. Full configuration files and analysis logs are deposited in the PhenoRewire repository (https://github.com/ldellavedova/PhenoRewire).
